# Distinct subdivisions of human medial parietal cortex are recruited differentially for memory recall of places and people

**DOI:** 10.1101/554915

**Authors:** Edward H Silson, Adam Steel, Alexis Kidder, Adrian W Gilmore, Chris I Baker

**Affiliations:** Laboratory of Brain & Cognition, National Institute of Mental Health, Bethesda, MD, USA; Wellcome Centre for Integrative Neuroimaging, FMRIB, Nuffield Department of Clinical Neurosciences, University of Oxford, Oxford, OX3 9DU, UK

## Abstract

Human medial parietal cortex (MPC) is implicated in multiple cognitive processes including memory recall, visual scene processing and navigation. It is also considered a core component of the default mode network. Here, we combine fMRI data across three independent experiments to demonstrate distinct subdivisions of MPC that are selectively recruited during memory recall of either specific places or specific people. First, distinct regions of MPC were identified on the basis of differential functional connectivity with medial and lateral regions of anterior ventral temporal cortex (VTC). Second, these same medial regions exhibited differential responses to the visual presentation of different stimulus categories, with clear preferences for scenes and faces, respectively. Third, and most critically, these regions were selectively recruited during either place or people memory recall. These subdivisions also showed a striking relationship with ventral and dorsal divisions of the default mode network. Taken together, these data reveal distinct subdivisions within MPC for the recall of places and people and moreover, suggest that the organizing principle defining the medial-lateral axis of VTC is reflected in MPC, but in the context of memory recall.

## Main text

Human medial parietal cortex (MPC), a core component of the default mode network (DMN)^1^ is associated with a diverse set of cognitive functions, including memory recall^2–5^ visual scene perception^6–8^, scene construction^9^, processing of spatial and other contextual associations^10^, navigation^11^, and future thinking^12^. Given such diverse recruitment of MPC across cognitive domains historically considered largely independent (e.g. visual processing, memory), it is not surprising that there appears little consensus in the literature as to the functional organization of MPC^13^. For example, MPC, along with ventral medial prefrontal cortex is thought to constitute the DMN core that flexibly integrates information between ventral and dorsal subnetworks of the DMN^1^. In contrast, others have utilized task-based^13^ and/or functional connectivity analyses^14–16^ to conceptualize MPC as fractionated beyond a simple core region but stopped short of describing a clear organizational structure. Thus, identifying a functional organization that provides synthesis across MPC is an important goal spanning multiple research fields.

Previously, our group^7^ and others^8,17^ have begun to elucidate the organization of MPC by demonstrating a strong functional link with anterior ventral temporal cortex (VTC). Specifically, a small region of MPC directly anterior of scene-selective medial place area^7^ (MPA) shows strong functional connectivity with anterior portions of scene-selective parahippocampal place area^18^ (aPPA), located in medial VTC. This connectivity-defined region overlaps, qualitatively, with regions of MPC engaged during memory recall^2–5^, suggesting that the ventral/posterior aspect of MPC may contain distinct areas biased toward scene processing for vision and memory, respectively^19^.

Whilst the previous functional link between MPC and VTC was based upon parcellating PPA along its posterior-anterior axis, the functional organization of VTC varies more dramatically along the orthogonal medial-lateral axis. Indeed, multiple functional dimensions are thought to be represented along this axis, including category preference^20,21^ (e.g. scenes, objects, tools and faces), eccentricity^22,23^ (e.g. peripheral, foveal), animacy^24^, and even real-world size^25^. Further, the mid-fusiform sulcus (MFS)^26^ has been identified as an anatomical landmark marking a transition point within each dimension (e.g. scene-selectivity medial of the MFS, face-selectivity lateral of the MFS). These robust functional differences across the medial-lateral axis of VTC arguably provide a stronger probe with which to investigate the organizational link between VTC and MPC.

To investigate the organization of MPC we conducted three independent fMRI experiments. First, we found that distinct subdivisions of MPC have preferential functional connectivity to anterior portions of medial and lateral VTC, respectfully. Second, these MPC subdivisions showed differential evoked responses to the presentation of different visual categories, with clear evidence for scene and face preferences. Third, and most critically, these subdivisions were selectively recruited during memory recall of either specific places (i.e. scenes) or specific people (i.e. faces). Finally, an independent whole-brain analysis of memory recall effects revealed an even finer division within MPC, with four identifiable regions showing an alternating (place/people) pattern of selective recruitment during memory recall.

Taken together, these findings provide converging evidence for a reflection of the functional organization of VTC in MPC. This organization was evident at rest, in response to visual stimulation, and most strikingly, during memory recall. The alternating pattern of responses throughout MPC provides a framework for understanding the broader functional organization of MPC. Collectively, these data support the notion that the functional organization defining the medial-lateral axis of VTC is reflected along the ventral/posterior-dorsal/anterior axis MPC, but in the context of memory retrieval.

## Results

### Subdivisions of MPC show preferential functional connectivity with medial and lateral portions of VTC

To determine whether the functional organization along the medial-lateral axis of VTC is reflected in MPC, we first utilized resting-state functional connectivity data (n=65). Six regions of interest (ROIs) were defined anatomically in each hemisphere that divided VTC along both the posterior-anterior and medial-lateral axes with respect to the MFS^26^, allowing us to characterize the connectivity profile between VTC and MPC more precisely **(Fig. 1a)**. A winner-take-all analysis **(see Methods)** revealed a ventral-posterior MPC region (referred to as MPC ventral, MPCv) that showed strongest connectivity with the anterior medial ROI and an adjacent dorsal-anterior region (referred to as MPC dorsal, MPCd) that showed strongest connectivity to the anterior lateral ROI **(Fig. 1b)**. Such a pattern of connectivity suggests a reflection of the functional organization defining the medial-lateral axis of VTC along the posterior/ventral-anterior/dorsal axis of MPC.

**Fig. 1:**
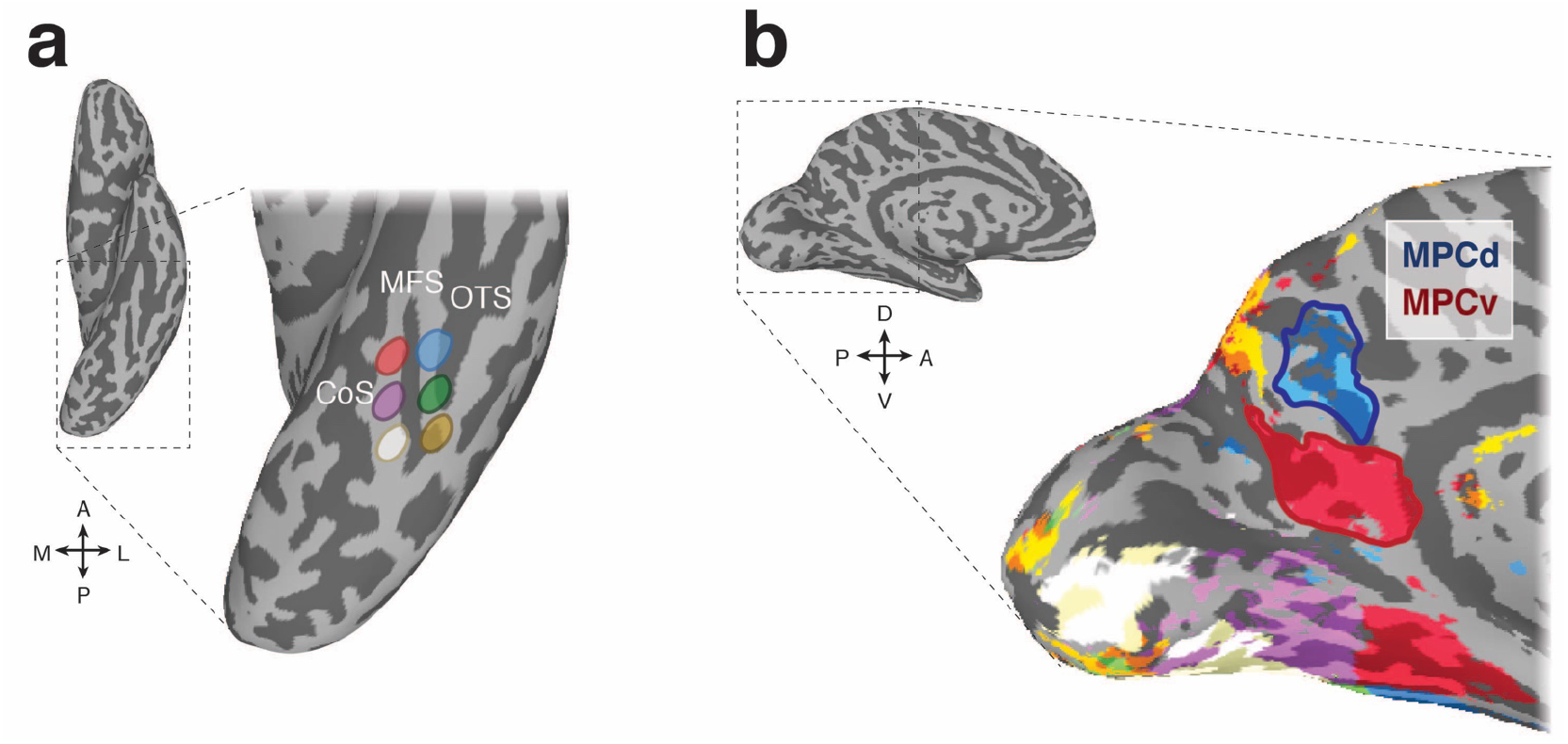
Resting-state functional connectivity seed regions and connectivity-defined regions of interest. **a**, A ventral view of the left hemisphere is shown with the ventral temporal cortex (VTC) highlighted with the dashed-black box, which is enlarged inset. Overlaid onto this enlarged surface are the six anatomically defined regions of interest that divide VTC along both the posterior-anterior and medial-lateral axes with respect to the mid-fusiform sulcus (MFS). The occipitotemporal sulcus (OTC) and collateral sulcus (CoS) are also labeled for reference. **b**. A medial view of the left hemisphere is shown with medial parietal cortex (MPC) highlighted by the dashed-black box, which is enlarged inset. Overlaid onto this enlarged surface is the result of the winner-take-all functional connectivity analysis. Colors on the brain correspond to the color of the anatomical ROIs in **a**. Within MPC, two separate regions are clearly visible. The ventral/posterior region (red-outline) is preferentially connected to anterior medial portions of VTC, whereas the dorsal/anterior region (blue-outline) is preferentially connected to anterior lateral portions of VTC. We define these resting-state ROIs as MPC ventral (MPCv) and MPC dorsal (MPCd), respectively.

### Subdivisions of MPC show differential responses to visually presented categories

Having identified distinct subdivisions of MPC based on differential functional connectivity with anterior portions of medial and lateral VTC, we next sought to determine whether these subdivisions would respond differentially to the visual presentation of different stimulus categories (Scenes, Faces, Buildings, Bodies, Objects and Scrambled Objects) **(see Methods)**. Differentiation on the basis of stimulus category would be reminiscent of the category-preference changes along the medial-lateral axis of VTC.

In a second independent group of participants (n=29), we calculated the mean response to each category (given as the *t*-value versus baseline) in both ROIs and hemispheres separately. Unlike category-selective regions of VTC (e.g. PPA, Fusiform Face Area, FFA^20^), which typically exhibit positive responses to the presentation of visual stimuli, we observed negative response magnitudes to all categories within both MPC subdivisions. Despite this general tendency for negative magnitudes, responses also appeared to differentiate on the basis of category, with scenes evoking the strongest response (i.e. least negative) in MPCv and faces evoking the strongest response in MPCd of both hemispheres **(Fig. 2)**.

**Fig. 2:**
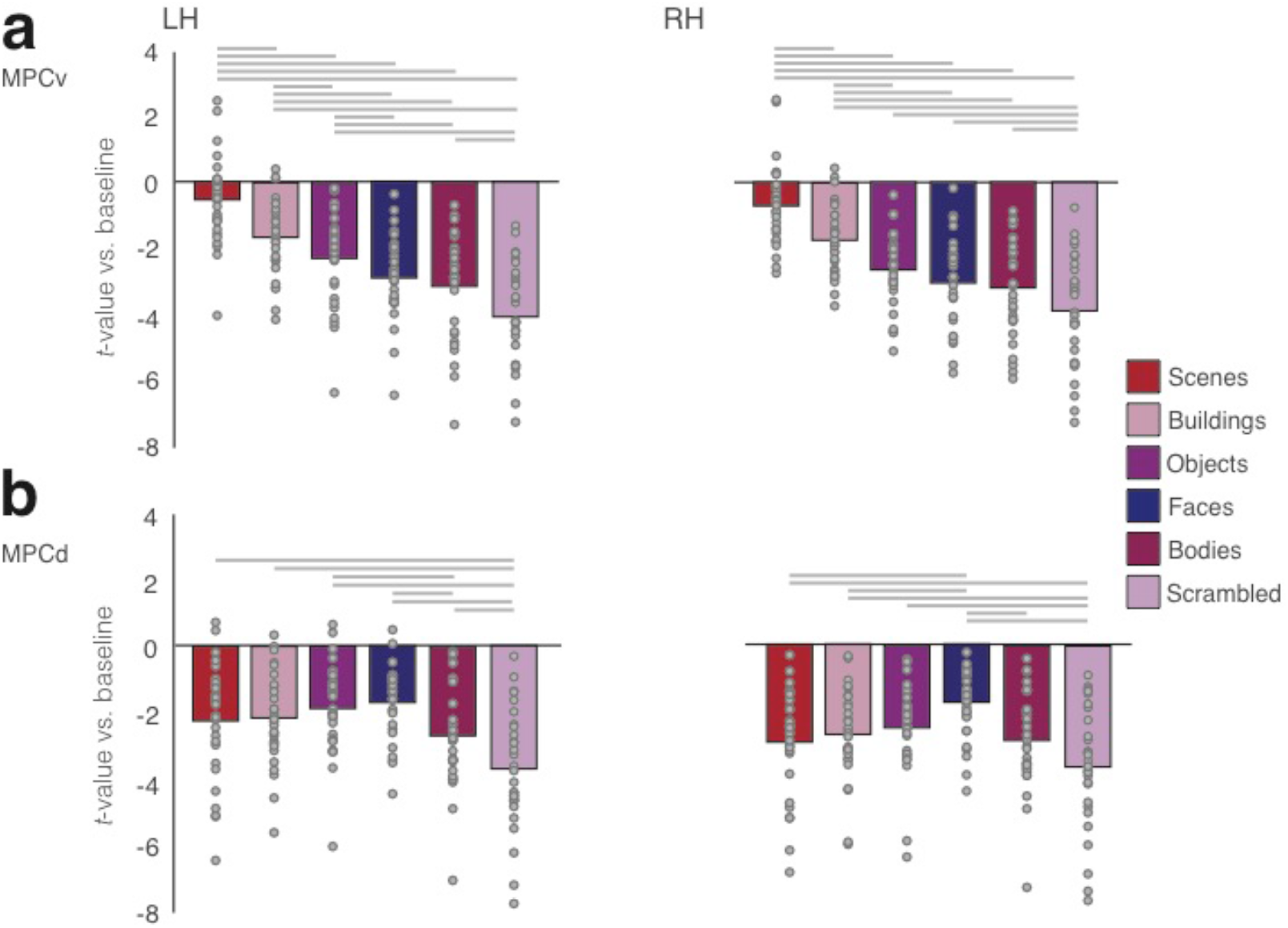
Negative responses in MPC to visually presented stimuli. **a**, Bars represent the mean response magnitude (given by the *t*-value versus baseline) for all six stimulus categories in MPCv of both hemispheres. These responses have been rank ordered from strongest (i.e. closest to baseline) to weakest. Single participant data points are shown for each category. Gray lines depict pairwise-comparisons that survived Bonferroni correction (*p*<0.01). The response to scenes was significantly different compared to all other categories in both hemispheres. **b**, Bars represent the mean response magnitude for all six stimulus categories in MPCd of both hemispheres. Bars are in the same order as in **a**, to highlight the different response profiles. Gray lines depict pairwise-comparisons that survived Bonferroni correction (*p*<0.01). The response to faces was significantly different from the majority of the other categories.

To explore these effects further, mean response magnitudes for each category were subjected to a one-way repeated measures Analysis of Variance (ANOVA) with Category (6 levels) as a within-participant factor. MPCv exhibited a significant main effect of Category in both the left (F_(5, 140)_=71.38, *p*<0.0001, partial η^2^=0.72) and right (F_(5, 140)_=49.46, *p*<0.0001, partial η^2^=0.64) hemispheres. Consistent with the stronger functional connectivity with medial VTC, the response to scenes was significantly different from other stimulus categories in both hemispheres (*t* > 5.34, *p*<0.001, in all cases, Bonferroni corrected) **(Fig. 2a)**.

MPCd also exhibited a significant effect of Category within both left (F_(5, 140)_=19.28, *p*<0.0001, partial η^2^=0.48) and right F_(5, 140)_=12.66, *p*<0.0001, partial η^2^=0.31) hemispheres. However, in this case the response to faces was only significantly different from Buildings and Scrambled Objects in the left hemisphere (*t* > 3.86, *p*<0.001, Bonferroni corrected; *p*>0.05, in all other cases), whereas the response to faces was significantly different from all categories except Objects (*p*>0.05) in the right hemisphere (*t* > 3.54, *p*<0.001, Bonferroni corrected) **(Fig. 2b)**. Collectively these results demonstrate a preference for scenes and faces within MPCv and MPCd, respectively. The overall pattern of negative responses evoked by visual stimuli is consistent with the widely-reported negative responses within MPC (and the broader DMN) when orienting to external stimuli^27^. However, motivated by the apparent scene and face preference within these regions and the fact that MPC is typically engaged positively during introspective tasks such as scene-construction from memory and mental imagery, we hypothesized that these MPC subdivisions would become differentially recruited during memory recall of either specific places (MPCv) or specific people (MPCd), respectively.

### Subdivisions of MPC differentially recruited during memory recall of specific places or specific people

To investigate this hypothesis, we conducted a memory recall experiment in a third independent group of participants (n=24). Participants performed six runs of a memory recall task, in which they were cued to recall from memory either specific places or specific people. Here, a simple 2 × 2 design was employed with two categories (Places, People) and two levels of familiarity (Famous, Personal) **(Fig. 3a)**.

**Fig. 3:**
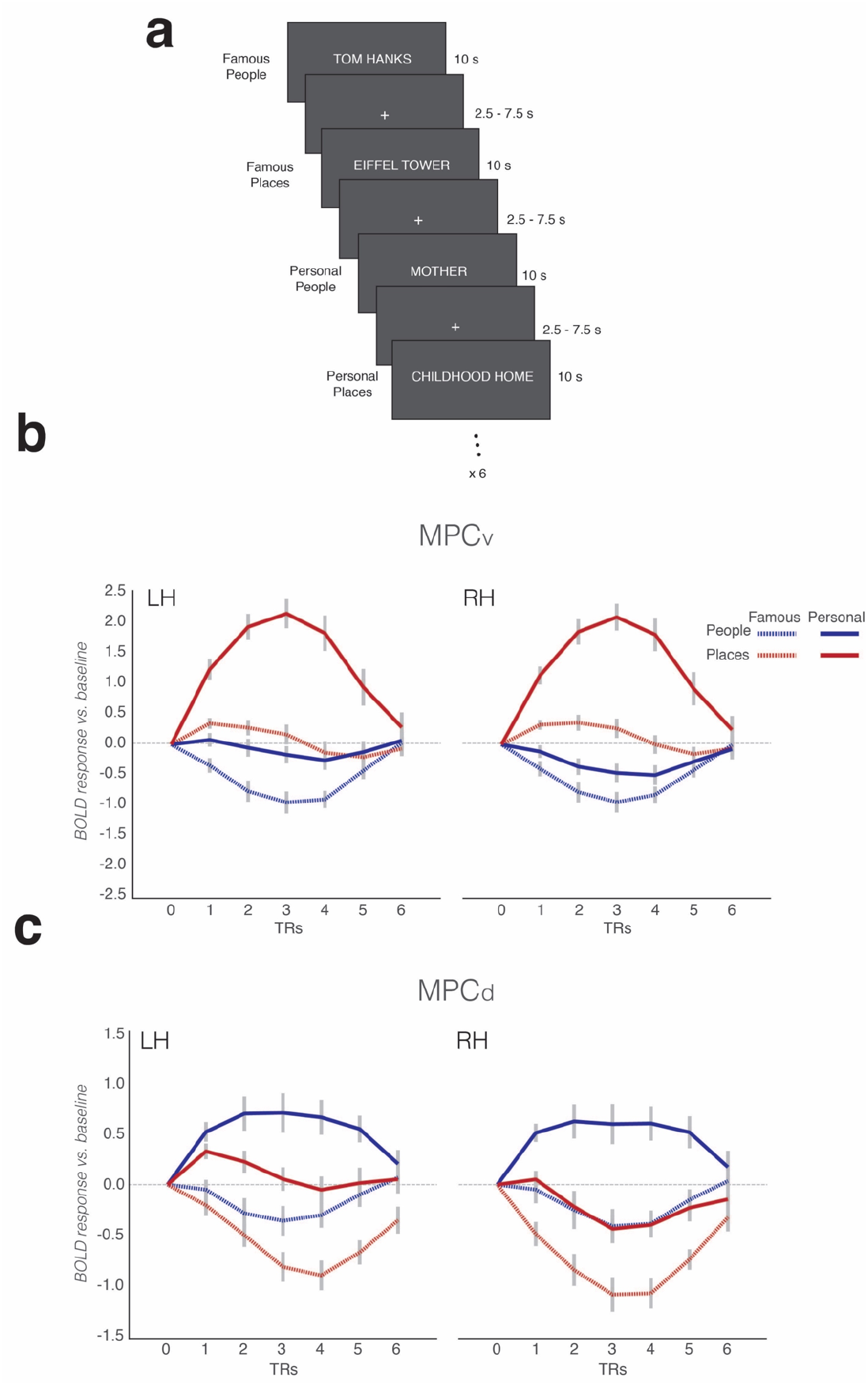
Memory task schematic and average BOLD responses to all conditions in MPC subdivisions. **a**, During the memory task, participants were given trial-wise instructions to recall from memory either famous people, famous places, personally familiar people or personally familiar places, respectively. Participants were asked to visualize the trial target from memory as vividly as possible for the duration of the trial (10 s). Trials were separated by a variable inter-trial-interval (2.5 −7.5 s). Participants completed 6 runs of the memory task. Each run contained 6 randomized trials from each condition. **b**, Average response curves from left and right MPCv (relative to baseline) are shown for all conditions in both hemispheres. Response curves were generated by first measuring the average response across the ROI for each trial (6 TR’s from trial onset) and then averaging across trails of the same condition. These responses were then averaged across participants and plotted for each condition separately (Famous people – dashed blue, Famous places – dashed red, Personal people – solid blue and Personal places – solid red). MPCv is maximally recruited during the recall of personal places. The patterns of response are very similar across hemispheres. **c**, Same as **b**, but for MPCd. In contrast to **b**, MPCd is maximally recruited during the recall of personal people. Again, this pattern is consistent across hemispheres. Response curves were normalized to begin at baseline (zero) for each trial separately. Gray-lines represent the standard error of the mean (sem) across participants for each condition and TR.

Subdivisions within MPC showed strikingly different response profiles during memory recall (derived by averaging the evoked responses across all trials per condition). Within MPCv, responses were maximally positive (relative to baseline) for the recall of personally familiar places, whereas responses during recall of famous people were maximally negative **(Fig. 3b)**. In contrast, responses within MPCd were maximally positive for the recall of personally familiar people and maximally negative during recall of famous places **(Fig. 3c)**. To quantify these responses, we calculated the mean contrast response (given by the *t*-value versus baseline) within each ROI to all conditions from the GLM analysis **(See Methods)**. These responses were then subjected to a three-way repeated measures ANOVA for each ROI separately, with Category (People, Places), Familiarity (Famous, Personal) and Hemisphere (Left, Right) as within-participant factors.

### MPCv selectively recruited during memory recall of specific places

Within MPCv, the main effects of Category (F_(1, 23)_=75.40 *p*=1.02^−8^, partial η^2^=0.76), Familiarity (F_(1, 23)_=128.61 *p*=6.78^−11^, partial η^2^=0.85) and Hemisphere (F_(1, 23)_=4.92 *p*=0.03, partial η^2^=0.17) were significant, reflecting on average greater responses for the recall of places over people, personal over famous stimuli and in the right compared to left hemisphere, respectively. However, these main effects were qualified by a significant three-way interaction (Category by Familiarity by Hemisphere: F_(1, 23)_=7.19 *p*=0.01, partial η^2^=0.24). This interaction reflects a larger familiarity difference (Personal > Famous) between categories (Place > People), in the right over left hemisphere. Further, we performed separate two two-way ANOVAs in each hemisphere separately with Category and Familiarity as factors. In both hemispheres, the Category by Familiarity interaction was significant (Left: F_(1, 23)_=31.19, *p*=0.00001, partial η^2^=0.56; Right: F_(1, 23)_=49.51, *p*=3.59^−7^, partial η^2^=0.68), reflecting a larger familiarity difference for the recall of places over people in both hemispheres **(Fig. 4a). (See Supplementary Material for full statistical breakdown)**.

**Fig. 4:**
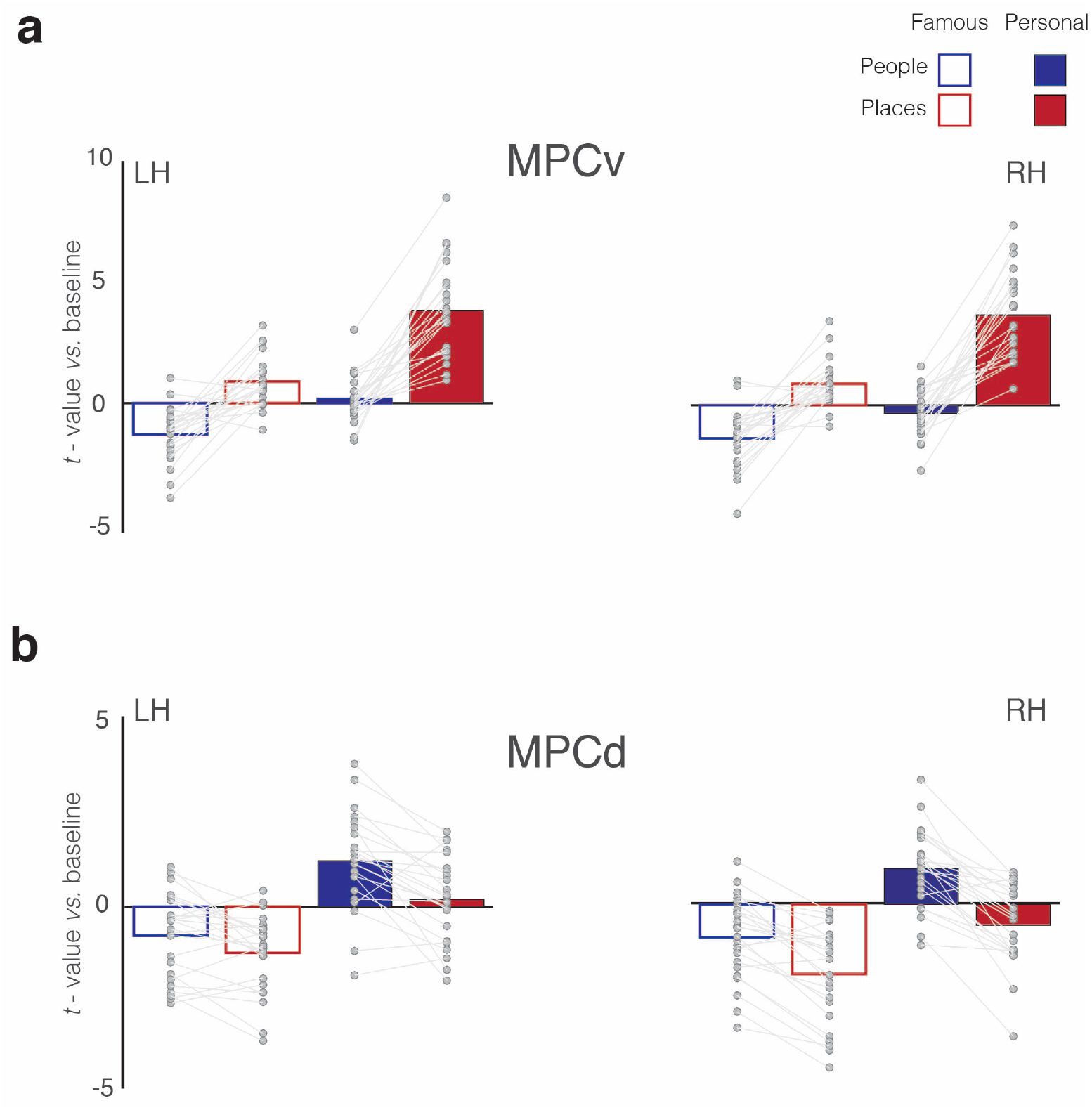
Magnitude of memory recall for all conditions in MPCv and MPCd. **a**, Bars represent the mean response magnitude for each condition (*t*-value versus baseline) in MPCv of both hemispheres (Famous people – blue open bars, Famous places – red open bars, Personal people – blue closed bars, Personal places – red closed bars). Data points for each participant are connected. In both hemispheres, MPCv is positively recruited during the recall of famous places and personal places, whereas responses during the recall of people (either famous or personal) are largely negative, reflecting a Category preference for places. MPCv also exhibits a familiarity effect and is maximally recruited during the recall of personal places, reflecting the effect of Familiarity. The interaction between Category and Familiarity is also evident. Indeed, there is a larger category difference (places-people) in the personal over famous conditions. **B**, Bars represent the mean response magnitude for each condition in MPCd of both hemispheres. Here, MPCd is only positively recruited during recall of personal people, reflecting both a Category preference for people and a Familiarity effect. The interaction between Category and Familiarity is also evident: there is a larger category difference (places-people) in the personal over famous conditions.

### MPCd selectively recruited during memory recall of specific people

Within MPCd, the main effects of Category (F_(1, 23)_=47.53 *p*=4.98^−7^, partial η^2^=0.67), Familiarity (F_(1, 23)_=82.33 *p*=4.62^−9^, partial η^2^=0.78) and Hemisphere (F_(1, 23)_=10.70 *p*=0.003, partial η^2^=0.32) were again significant, reflecting on average greater responses for the recall of people over places, personal over famous stimuli and in the right compared to left hemisphere, respectively. Whilst, we did not observe a significant threeway interaction, several significant two-way interactions were observed. Importantly, the Category by Familiarity interaction (F_(1, 23)_=7.89 *p*=0.01, partial η^2^=0.25), was significant, which reflects a larger familiarity difference for the recall of people over places with no clear difference between hemispheres **(Fig. 4b). (see Supplementary Material for full statistical breakdown)**.

### Consistent topography of memory recall effects within MPC

Both MPC subdivisions showed differential recall effects for places and people, respectively, coupled with an overall familiarity advantage. The topography of this differential recruitment during recall was strikingly consistent across individuals. Indeed, throughout MPC comparisons of the peak locations for the recall of personal places and personal people demonstrates a consistent shift along the ventral/posterior– dorsal/anterior axis. Across all participants and hemispheres, the peak response during recall of personal people was always anterior and dorsal to the peak response during recall of personal places **(Fig. 5)**.

**Fig. 5:**
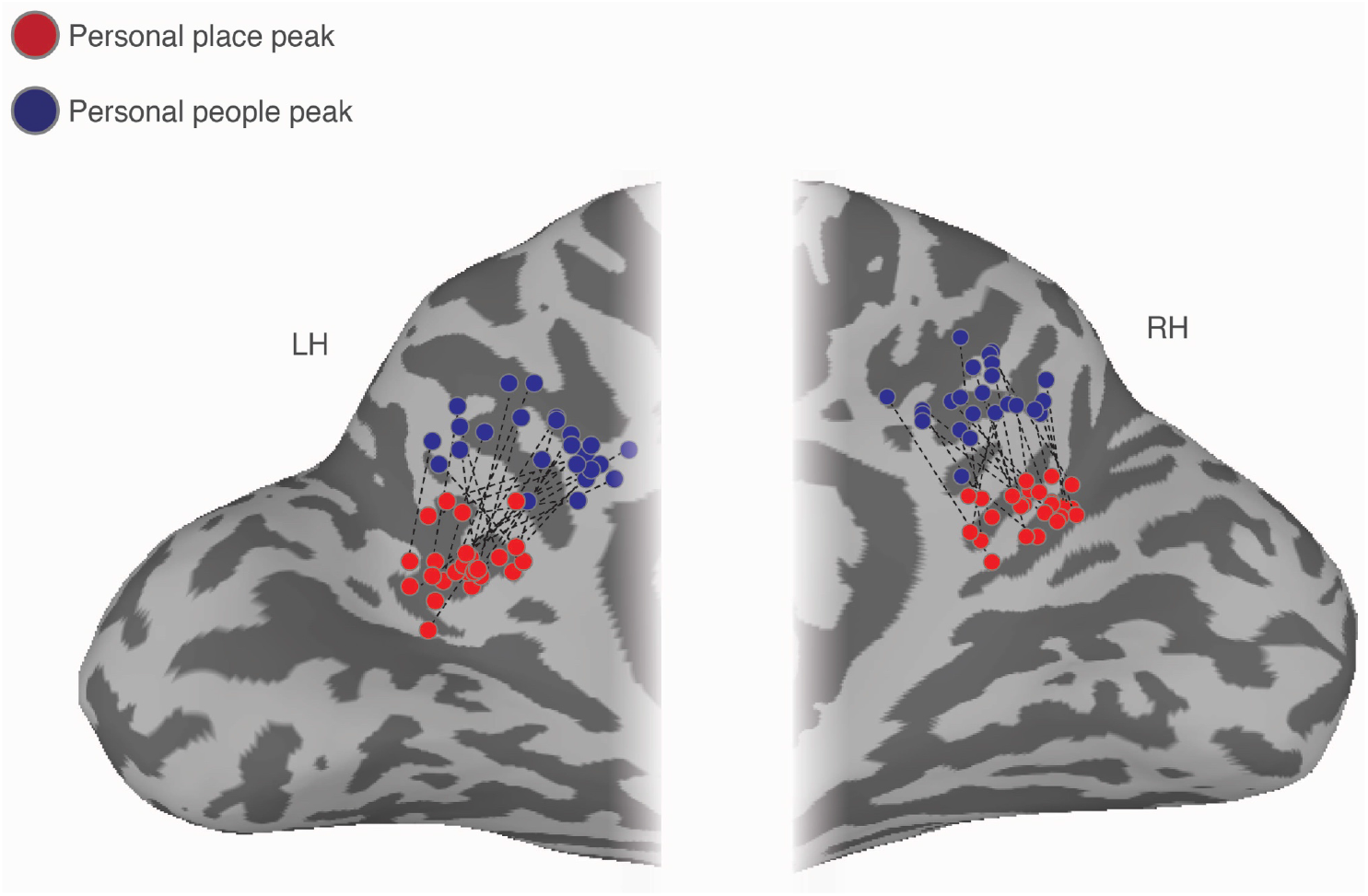
Ventral/posterior – dorsal/anterior shift in peak place memory and peak people memory. Enlarged partial views of the posterior medial portion of both the left and right hemispheres are shown. Overlaid onto these enlarged surfaces are the locations of the peak responses during recall of personal places (red dots) and recall of personal people (blue dots) for each participant. The corresponding peaks are connected for each participant with a dashed black line. Across participants, there is a consistent ventral/posterior – dorsal/anterior shift in the peak location of memory recall, such that the peak for recall of personal places is never posterior or ventral of the peak for recall of personal people.

### Alternating pattern of place and people memory recall throughout MPC

Having established that subdivisions of MPC are differentially recruited during memory recall, we next sought to determine whether areas outside of these initial ROIs showed similar effects. Accordingly, we performed a whole-brain Linear-Mixed-Effects (LME) modelling analysis to look for regions of the brain displaying main effects of Category (Places, People), Familiarity (Famous, Personal) and their interaction **(see Methods)**. At the whole-brain level, we did not observe any regions showing a significant interaction (at the selected statistical threshold), although significant responses to both main effects were present. The main effect of Category (collapsed across familiarity) revealed a complex pattern of differential recruitment throughout the brain. Most strikingly, along the ventral/posterior-dorsal/anterior axis of MPC, we observed an alternating pattern of memory recall: four adjacent subdivisions that alternated between being selectively recruited by the recall of places, then people, then places and finally people **(Fig. 6a)**. Notably, the first two subdivisions (refereed to here as ROIs 1 & 2) were largely equivalent to the connectivity-defined MPCv and MPCd **(see Supplementary Material for spatial overlap)**. Thus, this analysis not only confirmed the differential recruitment during memory recall of MPCv and MPCd, but also, revealed two anterior subdivisions within bilateral MPC that showed similar patterns of selective recruitment (ROIs 3 & 4). Strong memory recall for places was also present in aPPA in both hemispheres **(Fig. 6a)**. In contrast to the alternating pattern of place and people recall, the effect of familiarity (collapsed across category) manifested as an advantage for the recall of personally familiar over famous stimuli, irrespective of category within a relatively large swath of MPC **(Fig. 6b)**.

**Fig. 6:**
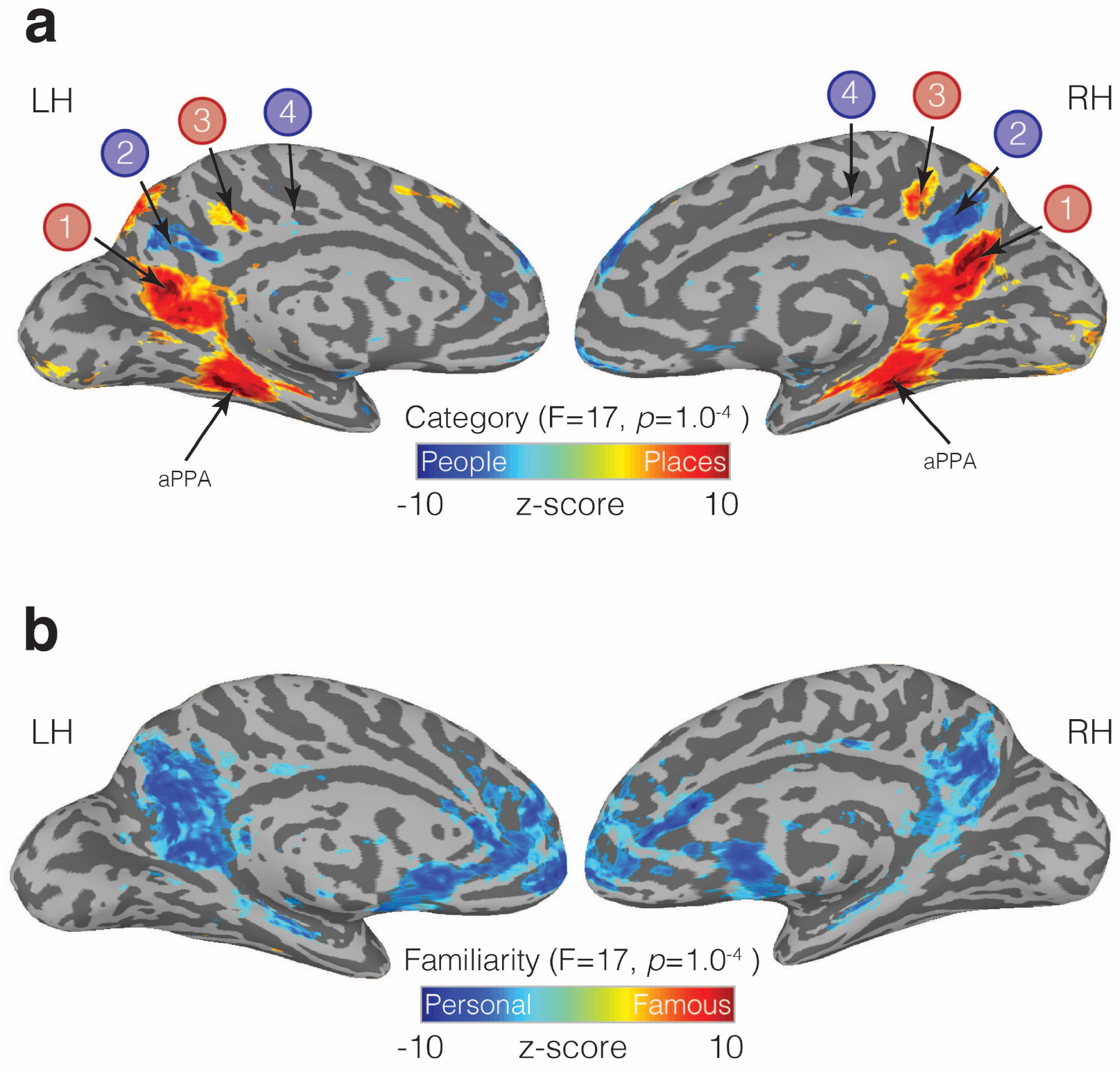
Familiarity and Category-selective recall in MPC. **a**, Medial views of both the left and right hemispheres of a representative participant are shown. Overlaid onto these surfaces is the main effect of Category from the linear-mixed-effects analysis (node-wise *p*=1.0^−4^, *q*=5.8^−4^). Cold colors represent regions of the brain more active during the recall of people (famous and personal), whereas hot colors represent regions of the brain more active during recall of places (famous and personal). An alternating pattern of memory recall is evident within MPC along the ventral/posterior – dorsal/anterior axis. ROIs 1 and 2, correspond largely to our initial resting-state ROIs (MPCv, MPCd), whereas the anterior pair of regions was not defined initially. We also observe significant place recall in aPPA, and some small clusters of significant people recall in anterior cingulate cortex. **b**, The same medial views are shown but overlaid is the main effect of Familiarity (node-wise *p*=1.0^−4^, *q*=5.8^−4^). Cold colors represent regions of the brain more active during the recall of personally familiar stimuli (places and people), whereas hot colors represent regions of the brain more active during the recall of famous stimuli (places or people). A large swath of MPC exhibits an overwhelming Familiarity effect with greater activity during recall of personal over famous stimuli. Familiarity effects were also present in the anterior cingulate cortex, insula and ventral medial prefrontal cortex.

To quantify the selective recruitment within each of these four MPC subdivisions in an independent manner, we implemented a split-half analysis. First, in each participant, we divided the six memory runs into odd and even datasets (3 runs each). Next, we performed the same LME analysis as above in each dataset separately **(see Methods)**. Between the odd and even splits, the topography and magnitude of the effect of category was robust and highly correlated across splits and hemispheres, respectively **(Fig. 7a)**. In order to determine estimates of effect size, we defined each MPC subdivision in one half the data (e.g. Odd) and sampled the responses to all conditions from the other half (e.g. Even). This process was then reversed, and the average computed. We observed a consistent and alternating pattern of selective recruitment throughout MPC. Recall of specific places selectively recruited ROIs 1 and 3, whereas recall of specific people selectively recruited ROIs 2 and 4. Consistent with our initial analyses, all four MPC subdivisions exhibited a familiarity advantage, which manifested as a selective enhancement in response for personally familiar items (Places = ROIs 1 and 3, People = ROIs, 2 and 4) **(see Supplementary Material for full statistical breakdown)**.

**Fig. 7:**
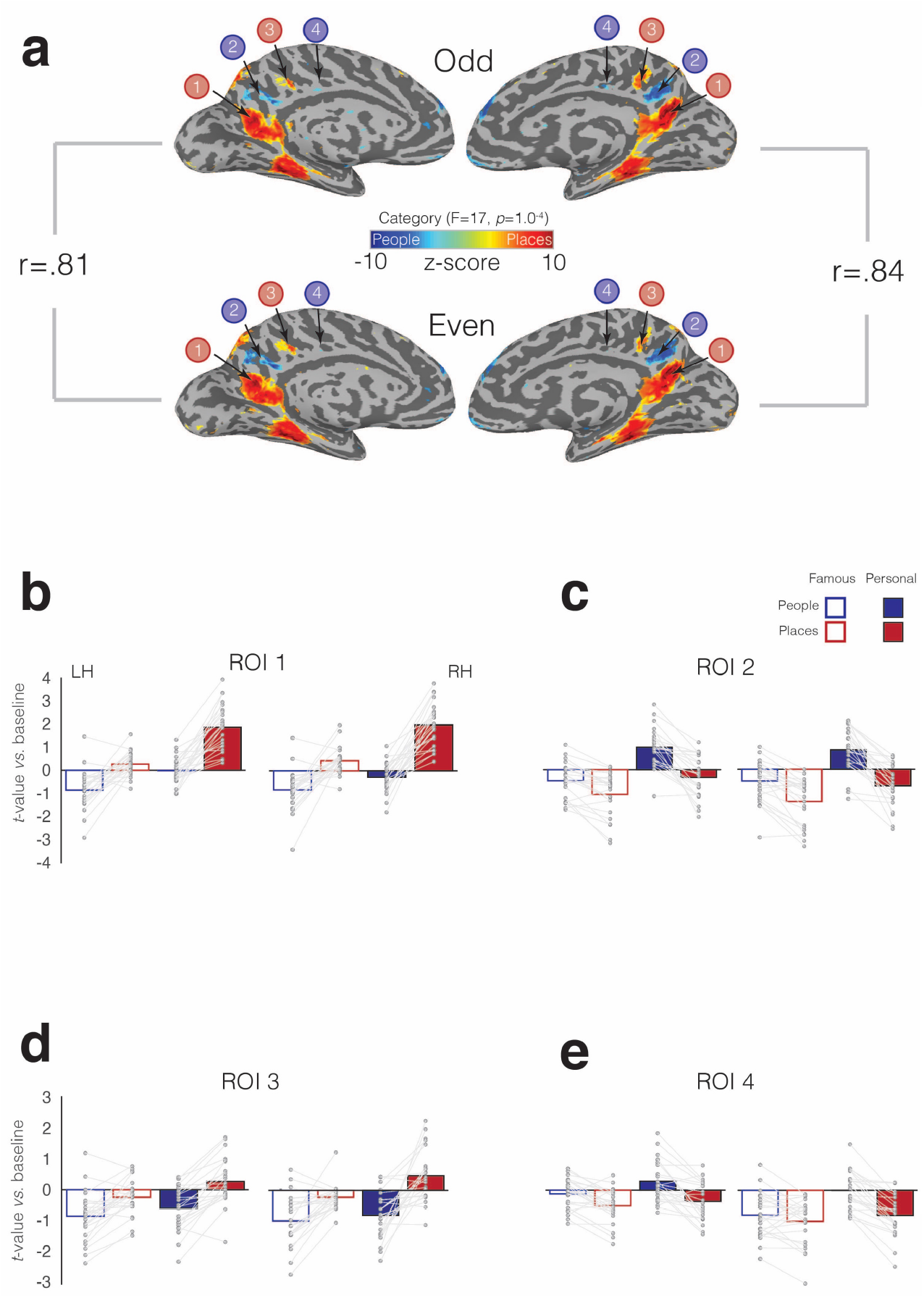
Split-half analysis and memory recall effects in four MPC subdivisions. **a**, The effect of Category (node-wise *p*=1.0^−4^, *q*=5.8^−4^) is overlaid onto medial views of both the left and right hemispheres for both independent halves of the data separately (Odd – top row, Even – bottom row). Cold colors represent regions of the brain more active during the recall of people, whereas hot colors represent regions of the brain more active during recall of places. Despite having half the amount of data, the alternating pattern of category-selectivity within MPC is present in both halves and hemispheres, respectively. The magnitude of the category effect (F-stat) was highly correlated across splits. The reported rho-values correspond to the correlation of the node-wise F-statistic for the effect of category in each hemisphere across the two splits. **b**, Bars represent the mean response magnitude for each condition (*t*-value versus baseline) from ROI 1. Single participant data points are shown and are connected for each participant. **c-e**, same as **a**, but for ROIs 2-4.

### Memory recall effects beyond MPC

Significant effects of place and people recall were also evident throughout the brain. In particular, the posterior angular gyrus, inferior temporal sulcus, and superior frontal regions were recruited during place recall, whereas the recall of people recruited the insula and anterior temporal regions, particularly in the right hemisphere **(See Supplementary Fig 1a)**. Advantages for the recall of personally familiar stimuli were present within anterior cingulate cortex and insula in both hemispheres **(Fig. 6b)**, as well as regions on the lateral surface, including superior-frontal, the superior-temporal sulcus, and angular gyrus/caudal inferior parietal lobule **(Supplementary Fig. 1b)**. In contrast, recall effects associated with famous over familiar stimuli were sparse and nonsignificant.

Memory recall effects were also observed within functionally-defined scene- and face-selective regions of VTC (i.e. PPA, FFA). Both regions were recruited during recall of items from their preferred category (i.e. greater response to place-specific memory in PPA, greater response to people-specific memory in FFA), although the magnitude of these memory effects were markedly weaker than within MPC (see **Supplementary Fig.2** and **Supplementary Material for full statistical breakdown**).

In addition to cortical ROIs, significant memory recall effects were also present in the hippocampus and amygdala bilaterally. The hippocampus showed an effect of category, with larger responses during recall of places over people, whilst also showing a strong familiarity advantage with larger responses for personally familiar over famous stimuli **(Supplementary Fig. 3a)**. In contrast, the amygdala showed only an effect of category, with larger responses during recall of people irrespective of the level of familiarity **(Supplementary Fig. 3b; see Supplementary Material for full statistical breakdown)**.

## Discussion

Across three independent fMRI experiments, we demonstrate subdivisions of MPC that exhibit i) differential functional connectivity to medial and lateral portions of anterior VTC, ii) show negative BOLD responses during visual perception with clear preferences for scenes and faces and iii) are differentially recruited during memory recall for either specific places or specific people. Taken together, these findings provide converging evidence that the functional organization defining the medial-lateral axis of VTC is reflected along the ventral/posterior-dorsal/anterior axis of MPC, but in the context of memory recall.

### The functional organization of medial parietal cortex

The selective recruitment of MPC during place or people memory is consistent with a diverse literature linking MPC with multiple memory processes^2–5^. Unlike previous neuropsychological^28–30^ and neuroimaging work^2–5^, which either lacked the spatial specificity to examine the heterogeneity of MPC or did not identify divisions of MPC precisely, we provide evidence for a distinct and heterogenous organization along the ventral/posterior-dorsal/anterior axis of MPC that appears to reflect the medial-lateral axis of VTC.

Initially, we focused on two MPC subdivisions, defined on the basis of preferential functional connectivity to medial and lateral portions of anterior VTC, respectively. These subdivisions exhibited different (albeit negative) responses to visually presented categories and were differentially recruited during memory recall. During the recall of specific places, MPCv showed strong positive evoked responses, which contrasted with negative evoked responses during the recall of specific people. In contrast, MPCd showed largely the opposite pattern - large positive evoked responses during recall of personally familiar people, but not during the recall of either famous people or places (either personal or famous). These data show a clear division between ventral/posterior and dorsal/anterior portions of MPC based on the content of the recalled memory and demonstrate a more heterogenous organization within MPC than previously reported^2–5, 28–30^. The cortical locations of MPCv/MPCd are consistent with previous observations of memory related activity^31–33^, but importantly, we also identify a second pair of anterior regions along the same axis, revealing a total of four subdivisions selectively recruited during memory recall for people or places in an alternating pattern.

A major contribution of the current work is the demonstration that the functional organization defining the medial-lateral axis of VTC is reflected along the ventral/posterior-dorsal/anterior axis of MPC. This complements and informs other recent parcellations of MPC^13–16^. For example, a recent meta-analysis attempted to divide a ventral portion of MPC (referred to as retrosplenial complex) on the basis of different fMRI task activations^13^. Although the majority of MPC was found to be recruited during memory tasks, a ventral/posterior region was more likely to be recruited during scene and navigation-related tasks, whilst more dorsal/anterior portions were more likely to be involved in theory-of-mind and social/emotional tasks. Similarly, a recent report^1^ separated social regions of MPC (i.e. those engaged with theory-of-mind tasks) from those involved with constructive memory or the formation of contextual associations^10^. These divisions align well with the differential memory recall effects reported here. The dorsal and ventral divisions also align well with recent studies that have attempted to map social network representations^34^ and representations of familiar scenes^35^, respectively. Our results provide a framework for understanding these previously-reported mnemonic effects by highlighting the apparent organizational link between the ventral/posterior-dorsal/anterior axis of MPC and the medial-lateral axis of VTC. Perhaps more importantly, these robust effects were evoked by the relatively simple task of recalling items that were cued by word stimuli only (or even merely perceiving presented stimuli), as opposed to performing more complex contextual association^10^, navigation^13^ or social judgment^33^ tasks. Consequently, this paves the way for future research to potentially address functional heterogeneity in MPC using tasks that target specific cognitive processes.

### Relating MPC to large-scale cortical networks

The role of MPC is often considered in the context of the DMN^1^. Although initially conceived as a singular entity^36^, the DMN has more recently been divided into two^38^ or three^37^ subnetworks. In the “ three network” framework, much of MPC and ventral medial prefrontal cortex act as a “core” that flexibly integrates information between the “dorsal” and “ventral” subnetworks^1,37^. Our results challenge this ‘core’ conceptualization by suggesting that MPC is fractionated along the same lines as these subnetworks: the dorsal component of the DMN, which overlapped regions recruited during people memory, is associated with social/semantic processing^38^ **(Fig. 8a)**, while, the ventral component, which is often referred to as a ‘scene construction’^9^ or ‘contextual association’^10,39^ network overlaps with regions recruited during place memory **(Fig. 8b)**.

**Fig. 8:**
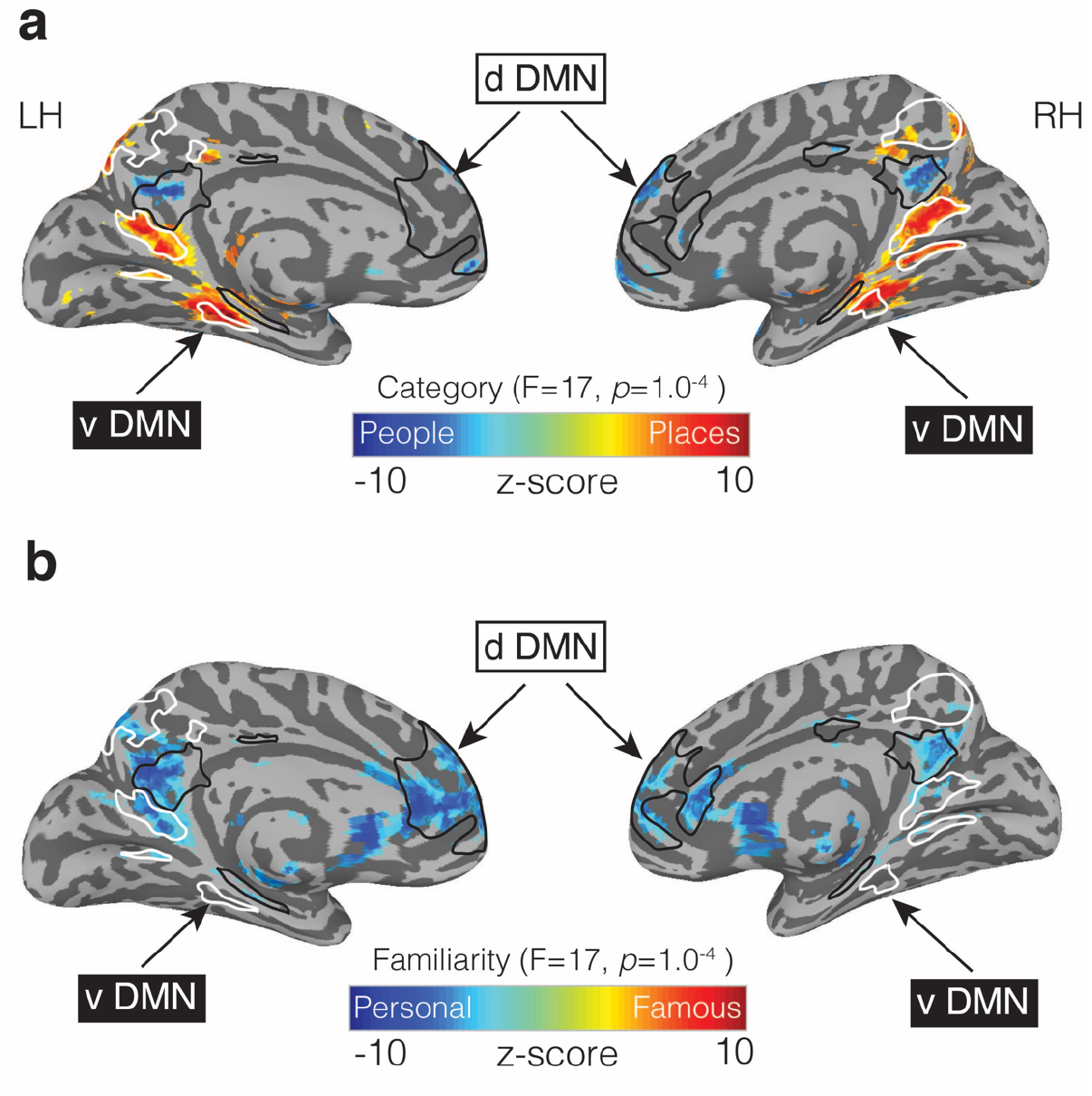
People and Place memory areas of MPC correspond to the dorsal and ventral DMN subnetworks. **a**, Medial views of both the left and right hemispheres are shown (TT n27 surface). Overlaid onto these surfaces is the effect of Category (*p*=1.0^−4^, *q*=5.8^−4^). Cold colors represent regions of the brain more active during the recall of people (famous and personal), whereas hot colors represent regions of the brain more active during recall of places (famous and personal). Masks of the dorsal DMN (dDMN) are outlined in black and show a striking spatial correspondence to regions more active during recall of people. In contrast, masks of the ventral DMN (vDMN) are outlined in white and correspond to regions more active during recall of places. DMN masks were taken from 38. **b**, The effect of Familiarity is overlaid onto the same surfaces (*p*=1.0^−4^, *q*=5.8^−4^). Cold colors represent regions of the brain more active during the recall of personally familiar stimuli (places and people), whereas hot colors represent regions of the brain more active during the recall of famous stimuli (places or people). Unlike in **a**, regions showing a Familiarity effect overlap with regions from both the dDMN and vDMN.

In the “ two network” framework^38^ the DMN is comprised of dorsal and ventral subnetworks without an integrative ‘core’. This framework is also consistent with recent work using highly-sampled participants to identify two parallel and interdigitated networks spanning cortex^15^. These two functional networks—both of which appeared to overlap with the canonical DMN—could be differentiated through functional connectivity, given sufficient data, but stopped short of describing their specific functional organization. In another sudy^40^, the authors noted the ventral MPC association with scene-related processing but did not comment on the more dorsal association with face processing. Our data provide a possible functional account of these networks by anchoring them to differential recruitment based on the content of perceived images as well as recalled memories, and moreover, demonstrate that such an interdigitated pattern of regions can be identified at the group-level given appropriate tasks.

The MPC regions we report also appear to overlap with the recently proposed “ posterior medial” (PM)^41^ memory system and the effects we observe are broadly consistent with the notion that the PM system is involved in recollecting episodic details, constructive uses of memory, and social cognition^41^. However, the present work suggests a differentiation within the PM system based on the recall of places and people, which was not discussed originally, suggesting that MPC’s role in memory recall does not fit neatly into a simple “ binary systems” model. More recently, a parietal memory network^4^ (PMN) has been identified, which includes regions within both lateral parietal cortex and MPC that are thought to be distinct from those within the DMN/PM^1, 41^ system. The memory effects we report overlap with DMN regions and appear close to, but separate from, those within the PMN. This adjacency reinforces the conceptual separation of processes associated with the PMN and the DMN/PM system, respectively.

### Functional correspondence between VTC and MPC

Multiple interrelated functional organizations are thought to be represented along the medial-lateral axis of VTC^20–25^. How the current data fit in the context of these organizations thus requires careful consideration.

Category-selectivity is one organizing principle thought to define the medial-lateral axis of VTC. Even at early stages of cortical development^21,23^, category-selective regions appear in stereotypical locations across individuals. This consistency across individuals and developmental time-frames^21^ supports the notion that categories are represented within discrete regions of VTC. The pattern of functional-connectivity, perceptual and memory recall effects we report are consistent with the presence of this motif within MPC. However, other functional dimensions are also thought to be represented along the medial-lateral axis of VTC, and the extent to which they interact and relate to one-another, and in-turn relate to the organization of MPC, is an open question^23,43^.

For example, eccentricity also varies systematically along the medial-lateral axis of VTC^22–23,42^. Indeed, these eccentricity representations are so highly correlated with category preference (e.g. PPA is peripherally biased^44^, whereas FFA is foveally biased^44^) that eccentricity has been suggested to form a prototypical organization onto which such selectivity later develops with visual experience^23,43^. It may be that the selective recruitment we observe in MPC during memory recall arises in a similar manner: That is, it is possible that the development of categorical-preference in VTC during perception drives the development of categorical-preference in MPC for memory. Alternatively, it is possible that the manner in which mnemonic representations of places and people are encoded and retrieved in MPC, differ in a way reminiscent of how the visual perception of places and people in VTC differ in terms of peripheral/foveal stimulation **(see Supplementary Fig. 4)**. The correspondence between MPC organization and the multiple dimensions thought to be represented across VTC, are key questions for future research. Importantly, these accounts are consistent with theories that suggest the organization of category representations in the brain are determined by the underlying structural (i.e. anatomical) and functional (e.g. perceptual) template^45^.

### Nature of responses within MPC during perception

The responses of MPCv/MPCd during perception share similarities with VTC but differ in important ways. For instance, responses in VTC during perception are characterized by larger evoked positive responses to stimuli of the preferred category^7,18,20^. In contrast, perceptual responses within MPCv/MPCd were best characterized by negative evoked responses that were attenuated, although not extinguished, by category-preference. Although the relationship between negative BOLD-signal changes and the underlying neural activity is an area of ongoing research^46–47^ and has been associated with inhibition in visual cortex^48^, this response is consistent with the widely-observed ‘task-negative’ activation of the larger DMN and the MPC component of it^1^.

## Conclusion

In this study we identified a consistent differentiation of regions within MPC, providing a new framework for understanding and investigating the functional organization of MPC and its role in memory retrieval. This differentiation was present at rest, in response to visually presented stimuli and finally through mnemonic recall. These data provide converging evidence that the functional organization defining the medial-lateral axis of VTC is reflected along the ventral/posterior-dorsal/anterior axis of MPC, but in the context of memory recall.

## Acknowledgments

We thank members of the Laboratory of Brain and Cognition for helpful comments on earlier version of the manuscript. This research was supported by the Intramural Research Program of the NIMH.

## Contributions

E.H.S, A.S & C.I.B jointly designed the study. E.H.S, A.S & A.K jointly collected the fMRI data. E.H.S & A.S jointly analyzed the fMRI data. E.H.S, A.S, A.K, A.W.G & C.I.B wrote the manuscript.

## Online Methods

### Participants

Participants for all experiments were recruited from the DC area and NIH community. All participants were right-handed with normal or corrected-to-normal vision and neurologically healthy. All participants gave written informed consent according to procedures approved by the NIH Institutional Review Board (protocol 93-M-0170, clinical trials # NCT00001360). Participants were monetarily compensated for their time.

#### Resting-state functional connectivity experiment

Sixty-five participants (40 female), mean age = 24.67 ± 3.2 years) completed the resting-state functional connectivity experiment.

#### Six category functional Localizer experiment

Twenty-nine participants (21 female, mean age = 24.2 years) completed the functional localizer experiment.

#### Memory experiment

Twenty-four participants (17 female, mean age = 24.2 years) completed the memory experiment.

### Stimuli and Tasks

#### Six category functional localizer experiment

Participants completed six functional localizer runs. During each run, color images from six stimulus categories (Scenes, Faces, Bodies, Buildings, Objects and Scrambled Objects) were presented at fixation (5×5° of visual angle) in 16 s blocks (20 images per block [300ms per image, 500ms blank]). Each category was presented twice per run, with the order of presentation counterbalanced across participants and runs. Participants responded via MRI compatible button box whenever the same image appeared sequentially. Stimuli for this and the other in-scanner tasks were presented using PsychoPy software^48^ (RRID:SCR_006571) from a Macbook Pro laptop (Apple Systems, Cupertino, CA).

#### Memory Experiment

Stimuli consisted of written names and words: 36 famous people, 36 famous places, 36 personally familiar people and 36 personally familiar places. The stimuli were provided by participants through a survey completed prior to the fMRI scan. Participants selected 36 known famous people and famous places from a list of 60 possible famous people (e.g., Tom Hanks, Angelina Jolie) and 92 possible famous places (e.g., Eiffel Tower, Times Square), and also provided the experimenters with the names of 36 people and 36 places that were personally familiar to them. Stimuli were presented in white 18-point Arial, all capital type against a black background.

During the task, on each trial participants were instructed to visualize the presented stimulus from memory as vividly as possible for the duration of the trial (10 s). Trials were separated by a variable inter-trial interval (2.5-7.5 s). In each of the six runs, there were six trials of each condition (famous people, famous places, personally familiar people and personally familiar places) presented in a randomized order, for a total of 24 trials per run (144 trials total).

#### Post scan questionnaire

After the scans were complete, participants completed a questionnaire in which they rated how vividly they were able to visualize from memory each stimulus presented during the memory runs. The stimuli were listed in the same order they appeared during the Memory Experiment and were rated on a 4-point Likert type scale (1 = not at all vivid; 4 = extremely vivid). If the participant could not visualize the stimulus at all while in the scanner, they checked a box on the questionnaire.

### Functional imaging parameters

Below we outline the imaging parameters for the three separate imaging experiments included in the current manuscript. All scans were performed on a 3.0T GE 750MRI scanner using a 32-channel head coil.

#### Resting-state functional connectivity

All functional images were acquired using a BOLD-contrast sensitive three-echo echo-planar sequence (ASSET acceleration factor = 2, TEs = =14.9, 28.4, 41.9 ms, flip-angle = 65°, bandwidth = 250.000 kHz, FOV = 24 × 24 cm, acquisition matrix = 64 × 64, resolution = 3.4 × 3.4 × 3.4 mm, slice gap = 0.3 mm, 34 slices per volume covering the whole brain). Respiratory and cardiac traces were recorded. Resting state scans lasted 21-minutes. The first 30 volumes were discarded to control for the state of arousal during the initial stages of data acquisition, leaving 20 minutes (600 volumes) for resting state functional connectivity analysis. This procedure has been used in other studies where long-duration resting state runs were collected.

#### Six category functional localizer

All functional images were acquiredusing a BOLD-contrast sensitive standard EPI sequence (TE=30 ms, TR=2 s, flip-angle = 65 degrees, FOV=192 mm, acquisition matrix = 64 × 64, resolution 2 × 2 × 2 mm, slice gap = 0.2mm, 37 slices covering the occipital and temporal lobes.

#### Memory experiment

All functional images were acquired using a BOLD-contrast sensitive three-echo echo-planar sequence (ASSET acceleration factor = 2, TEs = 12.5, 27.7, and 42.9 ms, flip angle = 75°, 64 × 64 matrix, in-plane resolution = 3.2 × 3.2 mm, slice thickness = 3.5 mm). Repetition times (TRs) and acquired slices varied across different task conditions to be consistent with relevant prior work for each task. The memory task used a 2500 ms TR, with 35 slices collected. All slices were collected obliquely and were manually aligned to the AC-PC axis.

### fMRI data preprocessing

Data were preprocessed using AFN^49^ (RRID: SCR_005927) for all experiments. Below we outline the preprocessing steps taken during each experiment.

#### Resting-state and memory experiments

The first 4 frames of each run were discarded to allow for T1 equilibration effects. Initial preprocessing steps for fMRI data were carried out on each echo separately. Slice-time correction was applied (3dTShift) and signal outliers were attenuated (3dDespike). Motion-correction parameters were estimated from the middle echo based on rigid-body registration of each volume to the first volume of the scan; these alignment parameters were then applied to the first and third echo. Data from all three acquired echoes were then registered to each participants’ T1 image and combined to remove additional thermal and physiological noise using multi-echo independent components analysis^50–51^. This procedure initially uses a weighted-average of the three echo times for each scan run to reduce thermal noise within each voxel. It subsequently performs a spatial ICA to separate time series components and uses the known properties of T_2_* signal decay to separate putatively neuronal BOLD components from putative noise components. This is accomplished by comparing each component to a model that assumes a temporal dependence in signal decay (i.e., that is “ BOLD-like”) and to a different model that assumes temporal independence (i.e., that is “ non-BOLD-like”). Components with a strong fit to the former and a poor fit to the latter model are retained for subsequent analysis (for further details, see^48^. This procedure was conducted using default options in AFNI’s tedana.py function. ME-ICA processed data from each scan were then aligned across runs for each participant.

#### Six category functional localizer experiment

All images were motion corrected to the first image of the first run (3dVolreg), after removal of the appropriate ‘dummy’ volumes (8) to allow stabilization of the magnetic field. Following motion correction, images were spatially smoothed (3dmerge) using a 5mm full-width-half-maximum smoothing kernel.

### fMRI data analysis

#### Resting-state functional connectivity winner-take-all analysis

Each participant’s resting-state functional connectivity time series was aligned to the standard surface using 3dVol2Surf. Six ROIs were defined that divided VTC along both the posterior-anterior and medial-lateral axes with respect to the mid-fusiform sulcus. For each participant, time series from these six ROIs were extracted, and the unique connectivity of each parcel to the rest of the brain was calculated using multiple-regression. The “ winning” parcel at each node was then determined by the maximum beta-value for each parcel (e.g. anterior medial VTC), and the selectivity index of the node was determined by subtracting the mean beta-values of all other parcels from the winning parcel (e.g. selectivity index = anterior medial VTC – (middle medial VTC + posterior medial VTC + anterior lateral VTC + middle lateral VTC + posterior lateral VTC).

#### Six category functional localizer analysis

A general linear model (GLM) approach was also used to analyze the functional localizer data. Specifically, a response model was built by convolving a standard gamma function with a 16 s square wave for each condition and compared against the activation time courses using Generalized Least Square (GLSQ) regression. Motion parameters and four polynomials accounting for slow drifts were included as regressors of no interest. To derive the response magnitude per condition, *t*-tests were performed between the condition-specific beta estimates (normalized by the grand mean of each voxel for each run) and baseline.

#### Memory analysis

Analyses were conducted using a general linear model (GLM) and the AFNI programs 3dDeconvolve and 3dREMLfit. The data at each time point were treated as the sum of all effects thought to be present at that time point and the time series was compared against GLSQ model fit with REML estimation of the temporal auto-correlation structure. Responses were modeled by convolving a standard gamma function with a 10 s square wave for each condition of interest (Famous People, Famous Places, Personal People and Personal Places). Estimated motion parameters were included as additional regressors of non-interest, and fourth-order polynomials were included to account for slow drifts in the MR signal over time. Significance was determined by comparing the beta estimates for each condition (normalized by the grand mean of each voxel for each run) against baseline.

#### Linear mixed effects analysis

To look at the whole brain memory effects we employed a linear-mixed-effects model (3dLME) in each hemisphere separately. The model comprised two factors: Category (Places, People) and Familiarity (Famous, Personal). At the whole brain-level, we did not observe any significant interactions, but both robust main effects were significant.

#### Split-half analysis

For each participant, the six memory runs were divided into Odd and Even splits (3 runs each). In each split, we performed the same LME. At the whole brain-level, we did not observe any significant interactions, but both robust main effects were significant. Throughout MPC we observed 4 ROIs that showed an alternating pattern of category-selective recall in both splits. To quantify these effects, we first defined each region within a split (e.g. Odd) and then sampled the data from the other half (e.g. Even). To avoid any potential bias in node selection, this process was reversed, and the average computed.

#### Sampling of data to the cortical surface

In each participant, the analyzed functional data were projected onto surface reconstructions of each individual participant’s hemispheres using the Surface Mapping with AFNI (SUMA) software. First, data were aligned to high-resolution anatomical scans (align_epi_anat.py). Once aligned, these data were projected onto the cortical surface (3dVol2Surf) and smoothed by a 2mm FWHM Gaussian kernel.

### Statistical Approach

Statistical analyses of both behavioral and functional data were performed using the SPSS software package (IBM). For all analyses we conducted repeated measures ANOVAs. When a significant three-way interaction was present, we performed two separate two-way ANOVAs to explore the nature of the interaction.

## Supplementary Material

**Supplementary Table 1:** Statistical analysis of behavioural responses collected directly after the memory experiment. Data are provided for both subjective vividness and the proportion of missed trials. Table includes F-values, degrees of freedom (*df*), *p*-values and estimates of effect size using partial eta squared. In each case, a two-way repeated measures ANOVA was conducted with Category (Places, People) and Familiarity (Famous, Personal) as within-participant factors. In the case of vividness ratings, both main effects of Category and Familiarity were significant, reflecting on average higher vividness ratings for the recall of people over places and for personal over famous stimuli. The significant Category by Familiarity interaction reflects a larger familiarity difference (Personal > Famous) during recall of places over people. For the proportion of missed trials, neither main effect was significant, but their interaction was. This interaction is driven by more missed trials for famous places than people, but fewer missed trials for personal scenes than people.

| 2-way ANOVA: Vividness |  |  |  |  |
| --- | --- | --- | --- | --- |
|  | *F-value* | *df* | *p-value* | *Partial eta squared* |
| Category | 23.35 | 1, 23 | 0.0007 | 0.50 |
| Familiarity | 53.93 | 1, 23 | 1.80<sup>-7</sup> | 0.70 |
| Category x Familiarity | 24.97 | 1, 23 | 0.0004 | 0.52 |
| 2-way ANOVA: Proportion Missed Trials |  |  |  |  |
| Category | 1.46 | 1, 23 | 0.23 | 0.06 |
| Familiarity | 1.64 | 1, 23 | 0.21 | 0.07 |
| Category x Familiarity | 7.26 | 1, 23 | 0.01 | 0.24 |

### Linear mixed effects analysis

To look at the whole brain memory effects we employed a linear-mixed-effects model (3dLME) in each hemisphere separately. The model comprised two factors: Category (Places, People) and Familiarity (Famous, Personal).

**Supplementary Figure 1:**
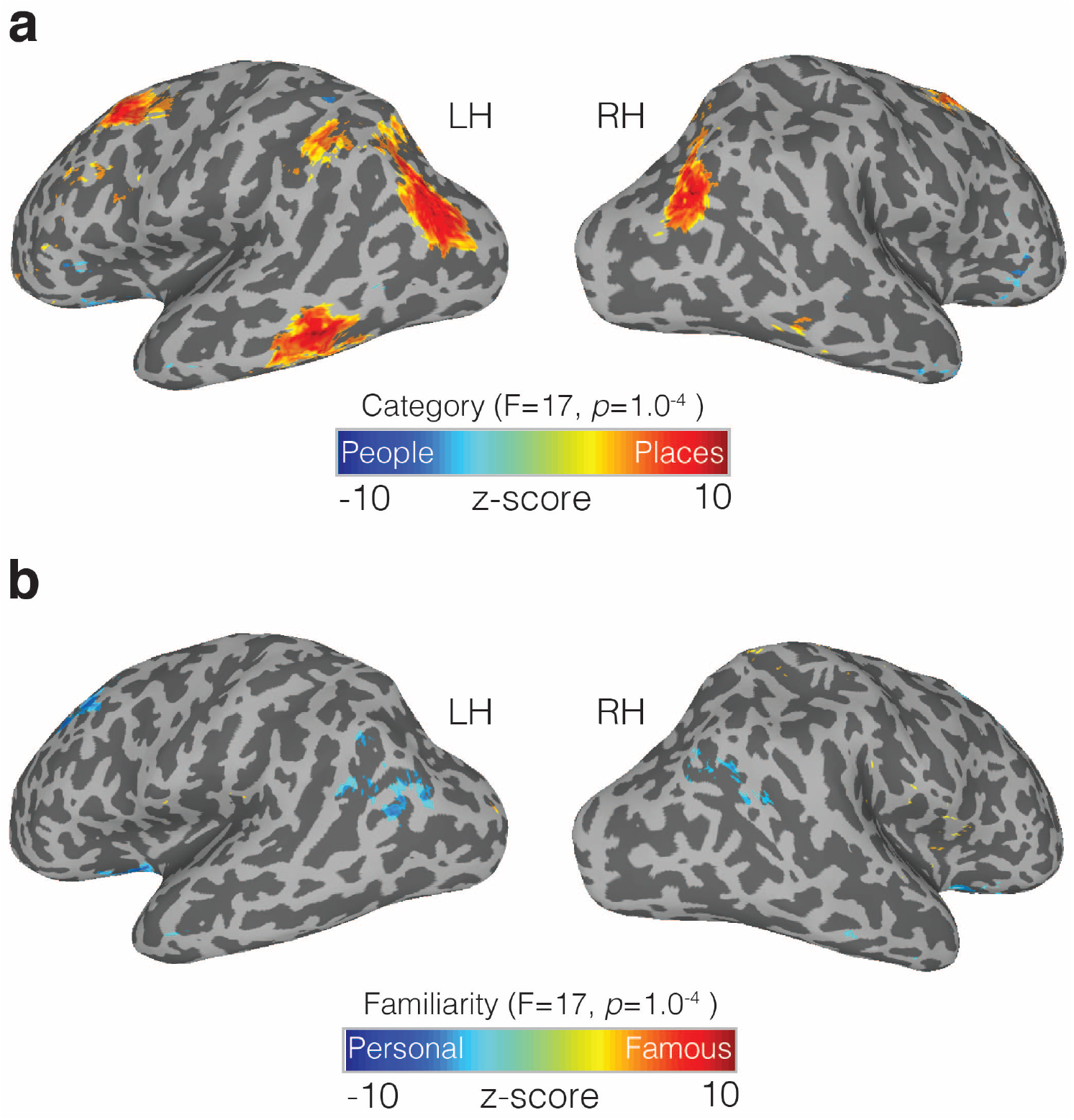
Memory recall effects on the lateral surface. **a**, Lateral views of the left and right hemispheres are shown. Overlaid onto these surfaces is the main effect of Category (*p*=1.0^−4^, *q*=5.8^−4^). Cold colors represent regions of the brain more active during the recall of people (famous and personal), whereas hot colors represent regions of the brain more active during recall of places (famous and personal). Significant responses during place recall were present in the posterior angular gyrus, inferior temporal sulcus and superior frontal regions. In contrast, significant responses to people recall were generally smaller in areal extent but nevertheless present in the insula and anterior temporal regions, particularly in the right hemisphere. **b**. Effect of Familiarity (*p*=1.0^−4^, *q*=5.8^−4^) overlaid onto the same surfaces. Cold colors represent regions of the brain more active during the recall of personally familiar stimuli (places and people), whereas hot colors represent regions of the brain more active during the recall of famous stimuli (places or people). Familiarity effects were present in superior-frontal regions, superior-temporal sulcus and the angular gyrus/caudal inferior parietal lobule.

### Statistical analysis of memory recall effects within ROIs

The same procedure was adopted when assessing memory effects from all ROIs, whether defined using resting-state (e.g. MPCv, MPCd), via the split-half analysis (ROIs 1-4), Category-selectivity (e.g. PPA, FFA) or via anatomical selection (e.g. Hippocampus, Amygdala). In each case, the mean response to each condition (given by the *t*-value versus baseline) was calculated for each participant, ROI and hemisphere separately. These values were then subjected to a three-way repeated measures ANOVA with Category (Places, People) Familiarity (Famous, Personal) and Hemisphere (Left, Right) as within-participant factors. If a significant three-way interaction was observed, we further explored the nature of this interaction with two-way ANOVAs in each hemisphere separately. Below are the full statistical breakdowns.

**Supplementary Table 2:** Statistical analysis of memory effects in MPCv and MPCd. Table includes Fvalues, degrees of freedom (*df*), *p*-values and estimates of effect size using partial eta squared. MPCv showed significant main effects of Category, Familiarity and Hemisphere. These were qualified by a significant three-way interaction, reflecting a larger familiarity difference (Personal > Famous) between categories (Places > People) in the right over left hemisphere. MPCd also showed significant main effects but did not show a significant three-way interaction. Importantly, however, MPCd did show the predicted Category by Familiarity interaction, which reflects a larger familiarity difference for the recall of people over places.

| 3-way ANOVA: MPCv |  |  |  |  |
| --- | --- | --- | --- | --- |
|  | *F*-value | *df* | *p*-value | *Partial eta squared* |
| Category | 75.40 | 1, 23 | 1.02<sup>-8</sup> | 0.76 |
| Familiarity | 123.86 | 1, 23 | 6.78<sup>-11</sup> | 0.85 |
| Hemisphere | 4.92 | 1, 23 | 0.03 | 0.17 |
| Category x Familiarity | 41.55 | 1, 23 | 0.000001 | 0.64 |
| Category x Hemisphere | 1.08 | 1, 23 | 0.31 | 0.04 |
| Familiarity x Hemisphere | 5.49 | 1, 23 | 0.03 | 0.19 |
| Category x Familiarity x Hemisphere | 7.19 | 1, 23 | 0.01 | 0.24 |
| 2-way ANOVA: Left Hemisphere |  |  |  |  |
| Category | 67.72 | 1, 23 | 2.63<sup>-8</sup> | 0.75 |
| Familiarity | 121.27 | 1, 23 | 1.20<sup>-10</sup> | 0.84 |
| Category x Familiarity | 31.19 | 1, 23 | 0.00001 | 0.56 |
| 2-way ANOVA: Right Hemisphere |  |  |  |  |
| Category | 78.25 | 1, 23 | 7.32<sup>-9</sup> | 0.77 |
| Familiarity | 111.74 | 1, 23 | 2.65<sup>-10</sup> | 0.83 |
| Category x Familiarity | 49.51 | 1, 23 | 3.59<sup>-7</sup> | 0.68 |
| 3-way ANOVA: MPCd |  |  |  |  |
| Category | 47.53 | 1, 23 | 4.98<sup>-7</sup> | 0.67 |
| Familiarity | 82.33 | 1, 23 | 4.62<sup>-9</sup> | 0.78 |
| Hemisphere | 10.70 | 1, 23 | 0.003 | 0.32 |
| Category x Familiarity | 3.79 | 1, 23 | 0.01 | 0.25 |
| Category x Hemisphere | 9.74 | 1, 23 | 0.005 | 0.28 |
| Familiarity x Hemisphere | 1.01 | 1, 23 | 0.32 | 0.42 |
| Category x Familiarity x Hemisphere | 0.22 | 1, 23 | 0.64 | 0.01 |

### Memory recall effects in the four MPC memory subdivisions

Of note, ROIs 1 and 2 although larger in areal extent shared considerable spatial overlap with our original resting-state definitions (ROI 1/MPCv proportional overlap, left=0.99, right=0.89; ROI 2/MPCd proportional overlap, left=0.55, right=0.67).

**Supplementary Table 3:** Statistical analysis of memory effects in ROI 1. Table includes Fvalues, degrees of freedom (*df*), *p*-values and estimates of effect size using partial eta squared. ROI 1 showed significant main effects of Category, Familiarity and Hemisphere. These were qualified by a significant three-way interaction, reflecting a larger familiarity difference (Personal > Famous) between categories (Places > People) in the right over left hemisphere.

| 3-way ANOVA: ROI 1 |  |  |  |  |
| --- | --- | --- | --- | --- |
|  | *F-value* | *df* | *p-value* | *Partial eta squared* |
| Category | 62.97 | 1, 23 | 4.92<sup>-8</sup> | 0.73 |
| Familiarity | 93.82 | 1, 23 | 1.39<sup>-9</sup> | 0.83 |
| Hemisphere | 0.07 | 1, 23 | 0.780 | 0.003 |
| Category x Familiarity | 32.39 | 1, 23 | 9.01<sup>-6</sup> | 0.58 |
| Category x Hemisphere | 12.42 | 1, 23 | 0.002 | 0.35 |
| Familiarity x Hemisphere | 10.05 | 1, 23 | 0.004 | 0.30 |
| Category x Familiarity x Hemisphere | 10.52 | 1, 23 | 0.004 | 0.31 |
| 2-way ANOVA: Left Hemisphere |  |  |  |  |
| Category | 60.24 | 1, 23 | 7.17<sup>-8</sup> | 0.72 |
| Familiarity | 80.37 | 1, 23 | 5.75<sup>-9</sup> | 0.78 |
| Category x Familiarity | 21.52 | 1, 23 | 0.0001 | 0.48 |
| 2-way ANOVA: Right Hemisphere |  |  |  |  |
| Category | 63.18 | 1, 23 | 4.78<sup>-8</sup> | 0.73 |
| Familiarity | 103.75 | 1, 23 | 5.40<sup>-10</sup> | 0.82 |
| Category x Familiarity | 43.16 | 1, 23 | 0.00001 | 0.68 |

**Supplementary Table 4:** Statistical analysis of memory effects in ROI 2. Table includes Fvalues, degrees of freedom (*df*), *p*-values and estimates of effect size using partial eta squared. ROI 2 showed significant main effects of Category, Familiarity and Hemisphere, but did not show a significant three-way interaction. Importantly, however, ROI 2 did show the predicted Category by Familiarity interaction, which reflects a larger familiarity difference for the recall of people over places.

| 3-way ANOVA: ROI 2 |  |  |  |  |
| --- | --- | --- | --- | --- |
|  | *F-value* | *df* | *p-value* | *Partial eta squared* |
| Category | 55.52 | 1, 23 | 1.42<sup>-7</sup> | 0.70 |
| Familiarity | 78.95 | 1, 23 | 6.76<sup>-9</sup> | 0.77 |
| Hemisphere | 6.42 | 1, 23 | 0.01 | 0.22 |
| Category x Familiarity | 17.19 | 1, 23 | 0.0003 | 0.42 |
| Category x Hemisphere | 5.45 | 1, 23 | 0.03 | 0.19 |
| Familiarity x Hemisphere | 1.19 | 1, 23 | 0.28 | 0.05 |
| Category x Familiarity x Hemisphere | 0.13 | 1, 23 | 0.72 | 0.006 |

**Supplementary Table 5:** Statistical analysis of memory effects in ROI 3. Table includes Fvalues, degrees of freedom (*df*), *p*-values and estimates of effect size using partial eta squared. ROI 3 showed significant main effects of Category, Familiarity, but not Hemisphere. Although ROI 3 did not show a significant threeway interaction, ROI 3 did show the predicted Category by Familiarity interaction, which reflects a larger familiarity difference for the recall of places over people.

| 3-way ANOVA: ROI 3 |  |  |  |  |
| --- | --- | --- | --- | --- |
|  | *F-value* | *df* | *p-value* | *Partial eta squared* |
| Category | 45.31 | 1, 23 | 7.25<sup>-7</sup> | 0.66 |
| Familiarity | 29.56 | 1, 23 | 0.00001 | 0.56 |
| Hemisphere | 0.24 | 1, 23 | 0.62 | 0.10 |
| Category x Familiarity | 6.93 | 1, 23 | 0.02 | 0.23 |
| Category x Hemisphere | 3.19 | 1, 23 | 0.08 | 0.12 |
| Familiarity x Hemisphere | 1.24 | 1, 23 | 0.27 | 0.05 |
| Category x Familiarity x Hemisphere | 3.47 | 1, 23 | 0.07 | 0.13 |

**Supplementary Table 6:** Statistical analysis of memory effects in ROI 4. Table includes Fvalues, degrees of freedom (*df*), *p*-values and estimates of effect size using partial eta squared. ROI 4 showed significant main effects of Category, Familiarity and Hemisphere. These were qualified by a significant three-way interaction, reflecting a larger familiarity difference (Personal > Famous) between categories (People > Places) in the right over left hemisphere.

| 3-way ANOVA: ROI 4 |  |  |  |  |
| --- | --- | --- | --- | --- |
|  | *F-value* | *df* | *p-value* | *Partial eta squared* |
| Category | 53.93 | 1, 23 | 1.80<sup>-7</sup> | 0.70 |
| Familiarity | 30.84 | 1, 23 | 0.00001 | 0.57 |
| Hemisphere | 30.44 | 1, 23 | 0.00001 | 0.57 |
| Category x Familiarity | 8.72 | 1, 23 | 0.007 | 0.27 |
| Category x Hemisphere | 0.06 | 1, 23 | 0.81 | 0.003 |
| Familiarity x Hemisphere | 4.61 | 1, 23 | 0.04 | 0.17 |
| Category x Familiarity x Hemisphere | 6.97 | 1, 23 | 0.02 | 0.23 |
| 2-way ANOVA: Left Hemisphere |  |  |  |  |
| Category | 27.12 | 1, 23 | 0.0002 | 0.54 |
| Familiarity | 10.60 | 1, 23 | 0.003 | 0.32 |
| Category x Familiarity | 2.84 | 1, 23 | 0.10 | 0.11 |
| 2-way ANOVA: Right Hemisphere |  |  |  |  |
| Category | 39.71 | 1, 23 | 0.00002 | 0.63 |
| Familiarity | 27.34 | 1, 23 | 0.0002 | 0.54 |
| Category x Familiarity | 12.38 | 1, 23 | 0.002 | 0.35 |

### Memory recall effects in category-selective VTC

Given that participants were recalling specific places (i.e. scenes) and specific people (i.e. faces) we also looked within scene (PPA) and face (FFA) selective regions of VTC for possible memory effects. These ROIs were defined at the group-level based on the contrast of Scenes > Faces (*p*=1.0^−4^).

**Supplementary Figure 2:**
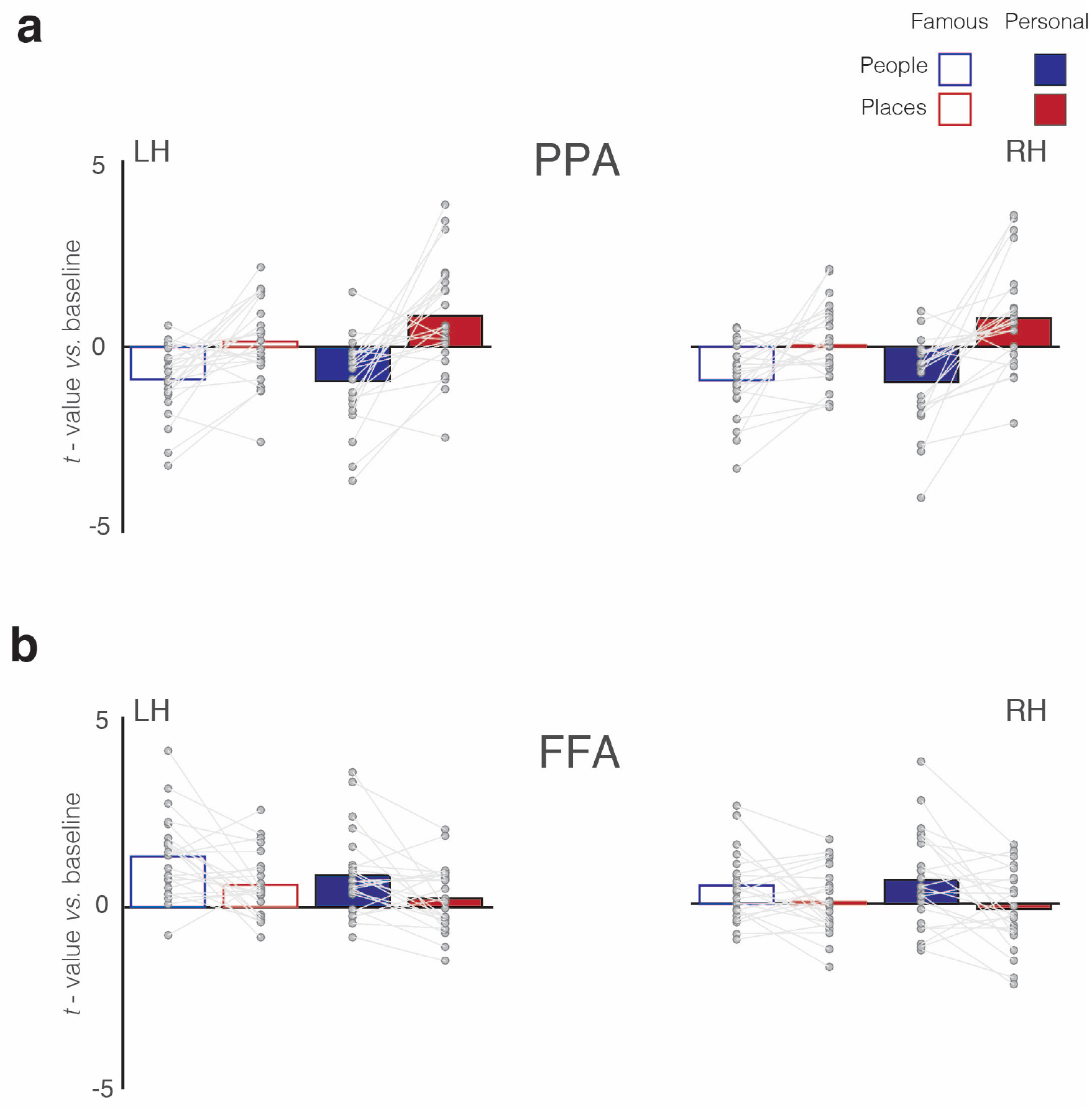
Memory recall effects in PPA and FFA. **a**, Bars represent the mean magnitude of response (*t*-value versus baseline) for all conditions in PPA of both the left and right hemispheres. Single participant data points and are shown and are connected for each participant. PPA is positively engaged during the recall of famous places and personal places, whereas responses during the recall of people (either famous or personal) are negative, reflecting a Category preference for places. PPA also exhibits a familiarity effect and is maximally engaged during the recall of personal places. The interaction between Category and Familiarity is also evident. Indeed, there is a larger category difference (places-people) in the personal over famous conditions. **b**, Same as **a**, but for FFA in both hemispheres. Unlike PPA, FFA is driven mostly by the recall of people but shows little to no effect of familiarity. Indeed, the response to famous and personal people is largely equivalent.

**Supplementary Table 7:**
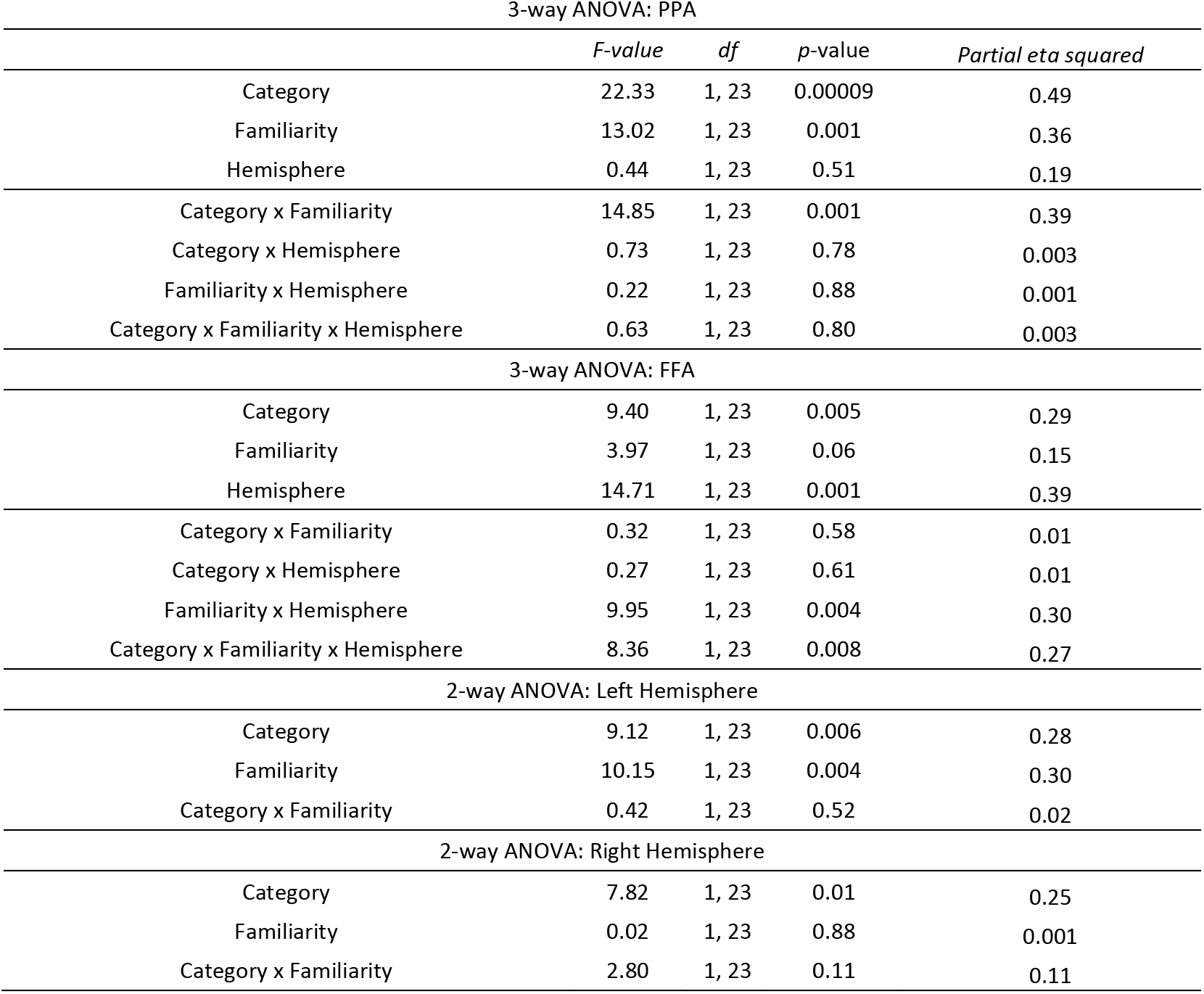
Statistical analysis of memory effects in PPA and FFA. Table includes Fvalues, degrees of freedom (*df*), *p*-values and estimates of effect size using partial eta squared. PPA showed significant main effects of Category and Familiarity, but not Hemisphere. PPA only showed a significant Category by Familiarity interaction, reflecting a larger familiarity difference (Personal > Famous) between categories (Places > People) with no clear difference between hemispheres. In contrast, FFA showed significant main effects of Category and Hemisphere, but not Familiarity. These were qualified by a significant three-way interaction, which reflects the presence of category and familiarity in the left hemisphere, but only the effect of category in the right hemisphere.

### Memory recall effects in Hippocampus and Amygdala

To better understand the memory recall effects observed in the Amygdala and Hippocampus detected at the group level, we sought to quantify these effects in individual participants using automatic parcellations of these structures in each participants’ native anatomical space.

**Supplementary Figure 3:**
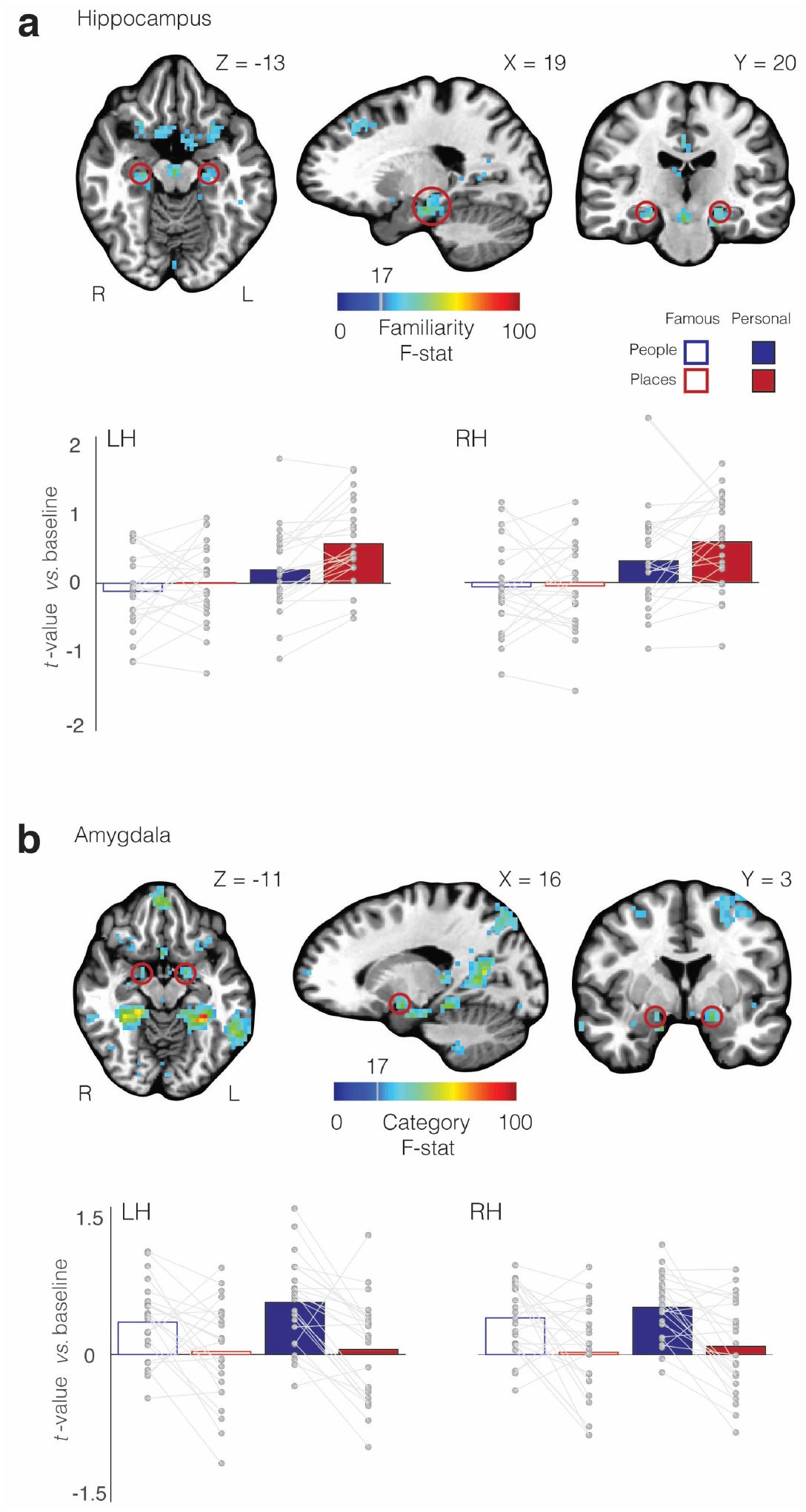
Memory recall effects in the Hippocampus and Amygdala. **a**, Axial, sagittal and coronal slices are shown. Overlaid onto these slices is the magnitude of the main effect of Familiarity. The red-circles highlight the approximate location of the Hippocampus in both hemispheres. Images are in radiological convention. Bars represent the mean magnitude of response to all conditions (versus baseline) in the Hippocampus of both the left and right hemispheres. Single participant data points and are shown and are connected for each participant. In both hemispheres, the Hippocampus shows larger positive responses on average during the recall of places over people, as well as, a familiarity effect with larger responses during recall of personal over famous stimuli. **b**, Axial, sagittal and coronal slices are shown. Overlaid onto these slices is the magnitude of the main effect of Category. The red-circles highlight the approximate location of the Amygdala in both hemispheres. Images are in radiological convention. Bars represent the mean magnitude of response to all conditions (versus baseline) in the Amygdala of both the left and right hemispheres. Unlike the Hippocampus, the Amygdala shows only the effect of Category with positive responses to the recall of famous and personal people, but little to no effect of Familiarity.

| 3-way ANOVA: Hippocampus |  |  |  |  |
| --- | --- | --- | --- | --- |
|  | *F-value* | *df* | *p-value* | *Partial eta squared* |
| Category | 5.51 | 1, 23 | 0.03 | 0.19 |
| Familiarity | 33.49 | 1, 23 | 0.000007 | 0.59 |
| Hemisphere | 0.13 | 1, 23 | 0.90 | 0.001 |
| Category x Familiarity | 3.27 | 1, 23 | 0.08 | 0.12 |
| Category x Hemisphere | 5.49 | 1, 23 | 0.03 | 0.19 |
| Familiarity x Hemisphere | 0.02 | 1, 23 | 0.88 | 0.001 |
| Category x Familiarity x Hemisphere | 0.11 | 1, 23 | 0.74 | 0.005 |
| 3-way ANOVA: Amygdala |  |  |  |  |
| Category | 20.77 | 1, 23 | 0.0001 | 0.47 |
| Familiarity | 2.91 | 1, 23 | 0.10 | 0.11 |
| Hemisphere | 0.16 | 1, 23 | 0.90 | 0.001 |
| Category x Familiarity | 0.87 | 1, 23 | 0.36 | 0.04 |
| Category x Hemisphere | 0.06 | 1, 23 | 0.80 | 0.003 |
| Familiarity x Hemisphere | 0.22 | 1, 23 | 0.64 | 0.01 |
| Category x Familiarity x Hemisphere | 2.41 | 1, 23 | 0.13 | 0.09 |

### Functional connectivity with foveal and peripheral early visual cortex

Given that category-preference and eccentricity are so highly correlated across the medial-lateral axis of VTC, we sought to test whether our initial MPC subdivisions would show stronger functional connectivity with foveal and peripheral portions of early visual cortex (EVC). Such a demonstration would provide further evidence for a functional link between VTC and MPC. Given the selective recruitment of MPCv during place recall, preference for scene stimuli and connectivity with medial VTC, we predicted stronger functional connectivity with peripheral over foveal portions of EVC. In contrast, given the selective recruitment of MPCd during people recall, preference for face stimuli and increased connectivity with medial VTC, we prediction the opposite pattern. To perform this analysis, we first defined foveal (0-4°) and peripheral (4-10°) regions of EVC in both hemispheres based on group retinotopic mapping data from a previous study from our group^43^. Next, we calculated the average functional connectivity between our two MPC subdivisions and these different eccentricity representations.

**Supplementary Figure 4:**
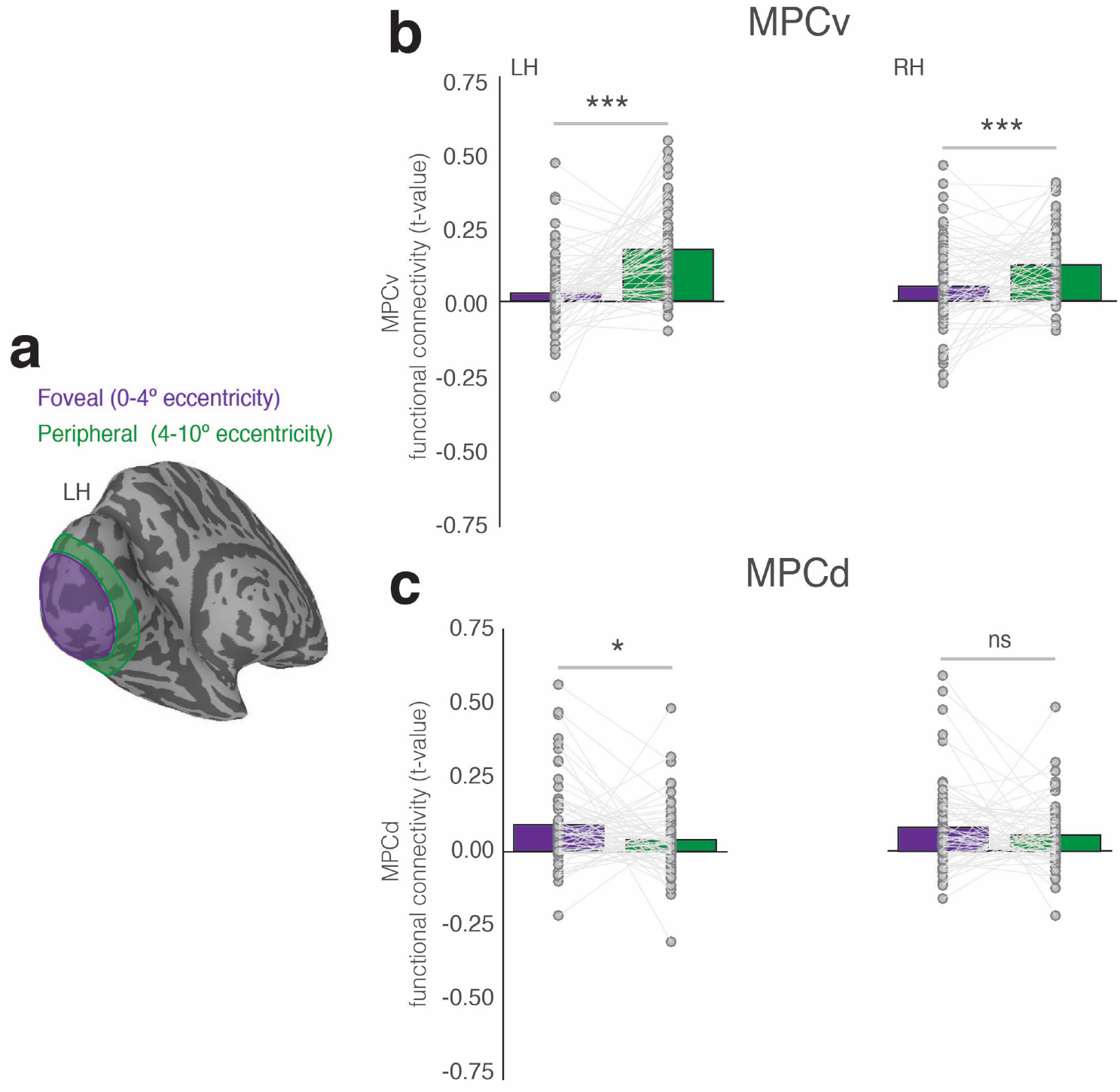
Functional connectivity between MPC subdivisions and foveal/peripheral early visual cortex. **a**, A posterior-medial view of the left hemisphere is shown with the foveal (purple) and peripheral (green) portions of early visual cortex highlighted. **b**, Bars represent the average functional connectivity between MPC places and both eccentricity representations in both hemispheres. Single participant data points and are shown and are connected for each participant. As predicted, MPC places shows on average stronger functional connectivity with peripheral over foveal portions of early visual cortex. **c**, same as **b**, but for MPC people. In contrast, MPC people shows on average stronger functional connectivity with foveal over peripheral portions of early visual cortex. ns = *p*>0.05, \**p*<0.05, \*\*\**p*<0.001.

**Supplementary Table 8:**
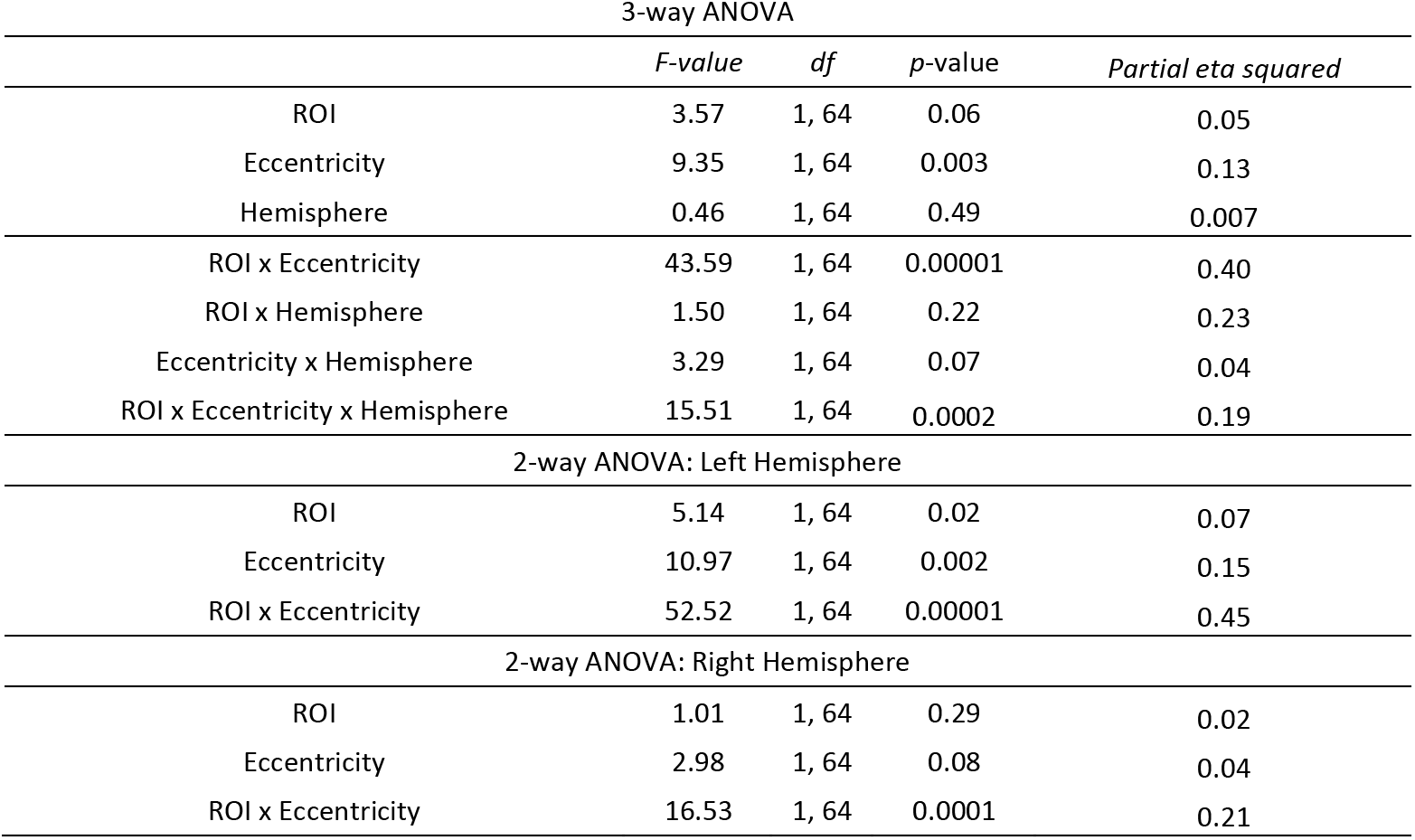
Statistical analysis of the resting-state functional connectivity between MPCv, MPCd and Foveal, Peripheral portions of early visual cortex (EVC). Table includes F-values, degrees of freedom (*df*), *p*-values and estimates of effect size using partial eta squared. The main effects of ROI and Hemisphere were not significant, but the main effect of Eccentricity was reflecting on average stringer connectivity with peripheral than foveal portions of EVC. These were qualified however by a significant three-way interaction, reflecting on average stronger connectivity between MPCv and peripheral EVC, but stronger connectivity between MPCd and foveal EVC in the left over right hemispheres.

| 3-way ANOVA |  |  |  |  |
| --- | --- | --- | --- | --- |
|  | *F-value* | *df* | *p-value* | *Partial eta squared* |
| ROI | 3.57 | 1, 64 | 0.06 | 0.05 |
| Eccentricity | 9.35 | 1, 64 | 0.003 | 0.13 |
| Hemisphere | 0.46 | 1, 64 | 0.49 | 0.007 |
| ROI x Eccentricity | 43.59 | 1, 64 | 0.00001 | 0.40 |
| ROI x Hemisphere | 1.50 | 1, 64 | 0.22 | 0.23 |
| Eccentricity x Hemisphere | 3.29 | 1, 64 | 0.07 | 0.04 |
| ROI x Eccentricity x Hemisphere | 15.51 | 1, 64 | 0.0002 | 0.19 |
| 2-way ANOVA: Left Hemisphere |  |  |  |  |
| ROI | 5.14 | 1, 64 | 0.02 | 0.07 |
| Eccentricity | 10.97 | 1, 64 | 0.002 | 0.15 |
| ROI x Eccentricity | 52.52 | 1, 64 | 0.00001 | 0.45 |
| 2-way ANOVA: Right Hemisphere |  |  |  |  |
| ROI | 1.01 | 1, 64 | 0.29 | 0.02 |
| Eccentricity | 2.98 | 1, 64 | 0.08 | 0.04 |
| ROI x Eccentricity | 16.53 | 1, 64 | 0.0001 | 0.21 |

